# Assessment and validation of enrichment and target capture approaches to improve *Mycobacterium tuberculosis* WGS from direct patient samples

**DOI:** 10.1101/2023.03.12.530724

**Authors:** B.C. Mann, K.R. Jacobson, Y. Ghebrekristos, R.M. Warren, M.R. Farhat

**Author notes:** Co-senior authors.

## Abstract

**Background:** Within-host *M. tuberculosis* (Mtb) diversity may capture antibiotic resistance or predict tuberculosis treatment failure and is best captured through sequencing directly from sputum. Here we compared three sample preprocessing steps for DNA decontamination and studied the yield of a new target enrichment protocol for optimal whole-genome sequencing (WGS) from direct patient samples.

**Design/Methods:** Mtb positive NALC - NaOH treated patient sputum sediments were pooled, and heat inactivated, split in replicates, and treated by either a wash, DNase I or Benzonase digestion. Levels of contaminating host DNA and target Mtb DNA was assessed by Quantitative PCR (qPCR), followed by WGS with and without custom dsDNA target enrichment.

**Results:** The pre-treatment sample has a high host-to-target ratio of DNA (6 168 ± 1 638 host copies/ng to 212,3 ± 59,4 Mtb copies/ng) that significantly decreased with all three treatments. Benzonase treatment resulted in the highest enrichment of Mtb DNA at 100-fold compared with control (3 422 ± 2 162 host copies/ng to 11 721 ± 7 096 Mtb copies/ng). The custom dsDNA probe panel successfully enriched libraries from as little as 0.45 pg of Mtb DNA (100 genome copies). Applied to direct sputum after benzonase treatment the dsDNA target enrichment panel increased the percent of sequencing reads mapping to the Mtb to 90,95% from 1,18 % compared with benzonase treatment without enrichment.

**Conclusion:** We demonstrate a low limit of detection (LoD) for a new custom dsDNA Mtb target enrichment panel that has a favourable cost profile. The results also demonstrate that pre-processing to remove contaminating extracellular DNA prior to cell lysis and DNA extraction improves the host-to-Mtb DNA ratio but is not adequate to support average coverage WGS without target capture.

## Introduction

*Mycobacterium tuberculosis* (Mtb) is the leading infectious pathogen killer globally (World Health Organisation (WHO), 2022). Although tuberculosis (TB) incidence has declined over the past decade, the global burden remains at more than 10 million people newly ill with the disease annually (Dheda *et al*., 2022). DNA sequencing advances over the past decade have enabled the study of Mtb’s genetic epidemiology, while also providing valuable information on within patient Mtb diversity, single nucleotide polymorphisms (SNPs) and other mutations which can be used to predict drug susceptibility (Brown et al., 2015; Goig *et al*., 2020a). Whole genome sequencing (WGS) of Mtb is still hindered by the long and cumbersome Mtb culturing process for DNA extraction. Culture can take weeks to months and has the additional limitation of potentially changing the population structure of the original sample due to selection of subpopulations more suited for growth in culture or stochastic purging from population bottlenecks (Nimmo *et al*., 2018, Goig *et al*., 2020a).

Sequencing the complete genome directly from clinical specimens would eliminate these issues. Several studies have demonstrated that sequencing from direct patient specimens is possible with varying levels of success (Brown *et al*., 2015; Votintseva *et al*., 2017; Doyle *et al*., 2018; Nimmo *et al*., 2018; Goig *et al*., 2020a; Soundararajan *et al*., 2020). The most successful direct from sputum sequencing (DSS) approach has used target specific RNA bait probes during library preparation. However, these probes still introduce several limitations, including low uniformity of coverage, the high proportion of duplicate reads generated as a result of PCR steps in both library preparation and enrichment, and technology costs ($110 – 168 per library and hybridisation reaction) (Bodi *et al*., 2013; Kozarewa *et al*., 2015). Probe based enrichment can also be limited by target capture specificity, making the approach sensitive to contamination and the ratio of contaminant to target DNA (Brown *et al*., 2015; Doyle *et al*., 2018; Nimmo *et al*., 2018; Goig *et al*., 2020a). A pre-processing step to decrease contaminant DNA burden can thus further boost enrichment of Mtb DNA for WGS (Goig *et al*., 2020a).

Prior studies have selectively lysed contaminating host and bacterial cells, followed by the depletion of contaminating DNA by enzymes such as DNase, either forgoing target enrichment or only using it in processing samples with low amounts of input mycobacterial DNA (Votintseva *et al*., 2017; Goig *et al*., 2020a). Although these studies were successful, this approach is highly dependent on Mtb bacillary load and proportions of contaminants including host cells and DNA as well as host microflora and their DNA (Votintseva et al., 2017). There is thus a need for a more consistently effective approach to contaminant depletion and Mtb target capture for DSS.

We postulated that sodium hydroxide (NaOH) decontamination followed by heat killing will lyse most contaminating host cells and non-Mtb bacteria, that Mtb cells are resilient to sodium hydroxide treatment and heat because of Mtb’s think and lipophilic cell wall, and thus after cell lysis the majority of remaining contaminating DNA will be extracellular and thus accessible to enzymatic degradation (Hasan *et al*., 2016; Islam *et al*., 2017). Previous studies have included DNase for depletion of extracellular contamination, but the direct effect on enrichment has not been systematically evaluated. Benzonase is an alternative enzyme for contamination removal that has not previously been evaluated (Votintseva et al., 2017; Goig *et al*., 2020a). Benzonase breaks down both ssDNA, dsDNA, RNA and DNA:RNA hybrids, while DNase targets primarily dsDNA with reduced specificity for ssDNA and DNA:RNA hybrids. Benzonase is thus an ideal candidate for extracellular DNA depletion that may provide a more efficient solution than the more frequently used DNase (Sutton *et al*., 1997; Liu *et al*., 2019; Amar *et al*., 2021).

In this study, we evaluate Benzonase and DNase head-to-head and compared their effect on enrichment and if enzymatic pre-treatment provides benefit to downstream target capture and enrichment. We also test the Twist (Twist Bioscience, USA) target capture system for Mtb target enrichment. The system is similar or more affordable ($110) than competing platforms and promises more uniform capture through the use of double-stranded DNA (dsDNA) biotin-labelled probes (Nagy-Szakal *et al*., 2022). Twist dsDNA probes have been used for Illumina sequencing from a range of starting materials, including respiratory specimens for SARS-CoV-2 WGS, and is hence a good candidate for sputum (Kim *et al*., 2021; Nagy-Szakal *et al*., 2022).

## Materials and methods

### Sample preparation

#### Limit of detection samples

Sputum sediment samples are expected to contain low amounts of total DNA, of which a small proportion is Mtb DNA. We generated a LoDseries of spiked samples using a H37Rv (ATCC 27294) reference strain to first assess the Twist kit’s ability to capture, and thus amplify, low amounts of Mtb target and prepare libraries from very low input concentrations (< 1 ng). Current manufacturer’s recommendations are to optimally start with 50 ng of input DNA but feedback from the manufacturers indicate that the kit can still successfully prepare libraries down to 1 ng. Liquid Middlebrook 7H9 medium supplemented with 0.2% glycerol, 0.25% Tween 80 (Sigma, USA.) and BD MGIT OADC enrichment supplement (BD, SA) was prepared, and 10 ml was dispensed into a filtered screwcap tissue culture flask 25 cm^2^ (Seperations, SA). The culture flask was then inoculated with the laboratory strain Mtb, H37Rv (ATCC 27294) and left to incubate for 2 weeks at 37 °C. Following incubation, the culture was transferred to a 15 ml falcon tube, centrifuged at 4000 rpm for 20 min and the supernatant was discarded. The pellet was then resuspended 200 μl lysis buffer prior to DNA extraction. Following extraction as described below, the DNA was diluted five times to generate samples consisting of approximately 10^5^, 10^4^, 10^3^, 10^2^ and 10 genomes/μl. This was estimated by considering the *Mtb* genome size of 4.4 Mb, expecting ~4.52 fg of gDNA to correspond to one Mtb genome. The starting sample had DNA concentration of 4.52 ng/μl or ~10^6^ H37Rv genomes and diluted 10-fold from there for each sample.

#### Sediment sample preparation and Pre-treatment to deplete contaminating host DNA

Routinely discarded Xpert positive sediments (NALC - NaOH decontaminated sputum samples – referred to as sediments) were collected from the National Health Laboratory Services Green Point, Western Cape, Cape Town, South Africa. Samples were frozen at – 20°C until further processing. Sediments were pooled to a final volume of 15 ml, heat inactivated for 1 hour at 80°C, removed from the biosafety level three facility and then aliquoted into 15 x 1 ml aliquots with intermittent mixing between each aliquot. The 15 aliquots were then split into groups of 3 replicates each per treatment condition namely a control, wash, benzonase treatment and DNase treatment group (Supplementary Figure 2). Sample treatment was subsequently processed as depicted in Figure 1.

**Figure 1:**
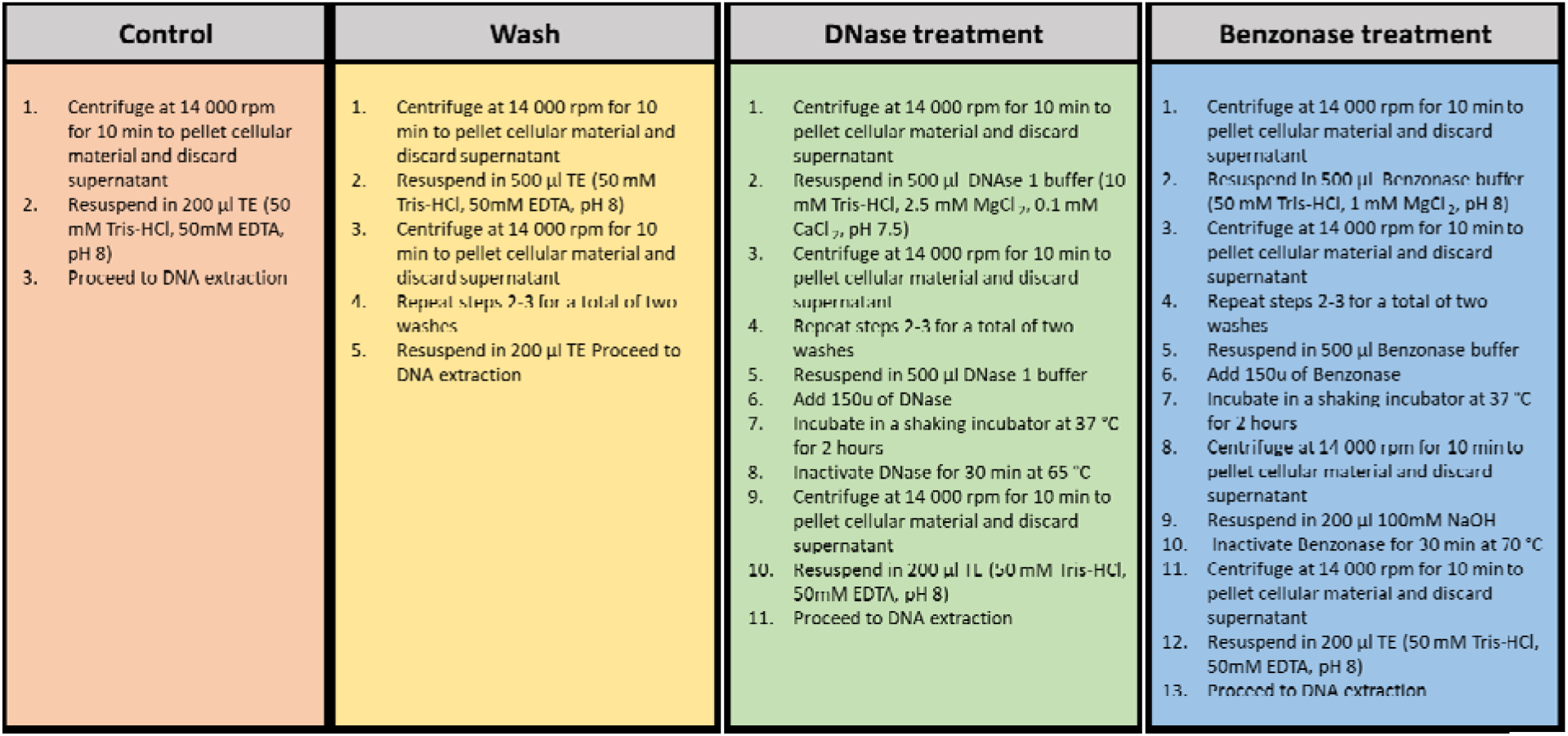
Summary of pre-treatment steps to deplete extracellular contaminating DNA prior to DNA extraction.

### DNA extraction

DNA was extracted using the Zymo - DNA Clean & Concentrator-25 (Zymo, USA) according to the manufacturer’s instructions (Epperson & Strong, 2020). Briefly, 50 μl of 100 mg/ml of lysozyme (Merck, Germany) was added to each sediment aliquot and incubated overnight at 37°C with gentle mixing. Thereafter, 50 μl of 2.5 mg/ml proteinase K (Merck, Germany) and 100 μl of 20 % SDS (Thermofisher Scientific, USA) was added and incubated at 65°C for 30 min. Binding buffer (800 μl) was added and gently mixed by inverting the tube until the sample solidified, then mixed vigorously by hand until the sample returned to a liquid state, and then gently mixed for 5 minutes. The solution was transferred to the column provided and centrifuged at 14 000 rpm for 30 seconds, this step was repeated until all of the solution had been loaded onto the column. The flow through was discarded after each centrifugation step. The column was then washed twice with 200 μl of wash buffer, by adding the wash buffer to the column and centrifugation at 14 000 rpm for 60 seconds. To elute the purified DNA, 50 μl of preheated elution buffer (10 mM Tris-HCl) is added to the column, incubated for 5 min and then centrifuged at 14 000 rpm for 30 seconds, the elution step was then repeated with an additional 50 μl aliquot of heated elution buffer to improve the final yield. The eluted DNA concentrations were quantified using the Qubit dsDNA HS Assay kit and the Qubit ssDNA Assay kit (Thermofisher Scientific, USA).

### qPCR

Luna Universal qPCR Master Mix (New England Biolabs, USA) and other reaction components were thawed at room temperature and gently vortexed. The final 20 μl volume for each reaction consisted of 10 μl 2X Luna Universal qPCR Master Mix (which contains deoxynucleotide triphosphate (dATP, dTTP, dCTP, and dGTP), MgCl^2^, Hot Start Taq DNA Polymerase, fluorescent dye and a passive reference dye), 0.5 μl forward primer (10 μM), 0.5 μl reverse primer (10 μM) (the human-specific target PTGER2 (primer pairs; hPTGER2f (5’- GCTGCTTCTCATTGTCTCGG −3’) and PTGER2r (5’- GCCAGGAGAATGAGGTGGTC - 3’) (Marotz et al., 2018), and the *Mtb* specific target Rv2341 (primer pairs; Rv2341-F (5’- GCCGCTCATGCTCCTTGGAT −3’) and Rv2341-F (5’- AGGTCGGTTCGCTGGTCTTG −3’) (Goig *et al*., 2020b), 5 μl nuclease free water and 4 μl of template (1 ng purified DNA). The “SYBR/FAM” channel of the CFX96 Touch Real-Time PCR Detection System was used for quantification (Biorad, USA). Cycling conditions were as follows: initial denaturation at 95°C for 1 min; 40 cycles consisting of 95°C for 15 s, 62°C for 30 s and 72°C for 30 s; and a final extension at 72°C for two minutes. All PCR amplification reactions were done in triplicate.

### qPCR analyses

Ct values were used to determine relative target abundance between samples or to measure absolute target quantities based on an acceptable standard curve obtained from a set of known dilutions. Standard curves were prepared at five concentrations by serial dilution, equating to input amounts of 10, 1, 0.1, 0.01 and 0.001 ng/per reaction of either *Mtb* or host DNA. The average Ct for each sample was taken and used along with the standard curve to calculate the approximate copy number of either *Mtb* or host DNA to ascertain the ratio of target to host DNA (*Mtb* DNA:host DNA). Copy number estimates were normalized per nanogram input DNA.

### Library preparation and target capture and enrichment

Twist target capture probes were custom designed for 1-fold coverage of the full H37Rv genome with duplicates removed, supplemented by 4-fold coverage of 95 known lineage single nucleotide variants (SNVs) (Freschi et al., 2021) and 1387 homoplasic SNVs and 60 INDELs in drug resistance regions (Supplementary folder 1 - Probe panel). Targeted mutations were merged within a 60bp region and tiled as a block 4-fold. Probes were manufactured by Twist Biosciences, San Francisco, USA, and shipped to Stellenbosch University, South Africa.

The limit of detection range as well as 1 sample from each treatment condition of the benzonase/DNase head-to-head comparison experiment were selected for Twist target capture and sequencing (Characteristics for the final pool of samples can be found in Supplementary Table 5). All 8 samples were subjected to library preparation using the Twist enzymatic fragmentation and universal adapter system as follows. Library preparation was done according to the manufacturer’s instructions with only two modifications. Considering the low input concentrations and based upon feedback from the manufacturer the adapter concentration was halved and the amplification cycles using the TWIST UDI primers during the library preparation step was increased from 8 to 10 cycles and the. The 8 prepared libraries were pooled for hybridisation and post capture amplification, the resulting library was run on a Miniseq sequencer (Illumina, USA), using the MiniSeq High Output Reagent Kit (150-cycles).

### Bioinformatics

Initial quality assessment was done using fastQC, version 0.11.9, followed by adapter trimming, quality filtering and per-read quality pruning using fastP, version 0.20.1 (Chen *et al*., 2018). Following quality control steps, reads were taxonomically classified using the metagenomic classification tool Kraken2 (version 2.0.8). A custom kraken2 database was built using GenBank/RefSeq sequences, and included all GenBank bacteria, fungi, plasmid, viral and phage sequences (complete, chromosome and scaffold). The RefSeq human genome along with additional *Mtb* genomes (CP011510_1, NC_000962_3, NC_002755_2, NC_009565_1, NC_017524_1, NZ_OW052188_1 and NZ_OW052302_1) were also added to the database (Wood *et al*., 2019). Reads classified as Mtb were extracted using the krakentools extract_kraken_reads.py script. Both filtered and unfiltered fastq files were aligned using bwa-mem2 (version 2.2.1) to the H37Rv reference genome AL123456 (Vasimuddin *et al*., 2019). Duplicates were removed using Picard tools and excluded in further downstream analyses. Alignment statistics, including number of reads, depth and breadth of coverage and GC-content were determined and visualised using Qualimap (version 2.2.2c). Outputs from Qualimap were used to generate plots of the normalized sequencing depth across the reference genome and any observed regions with deviations in coverage were visualised using Integrative Genomics Viewer (version 2.3.97) (GarcÃa-Alcalde *et al*., 2012; Thorvaldsdottir *et al*., 2013).

### Statistical analyses

Statistically significant differences between control and treatment groups were determined using the unpaired t-test with Welch’s correction, for both the DNA concentrations as determined by the ss and dsDNA Qubit assays, as well as for the copy number estimates based on qPCR results.

## Results

### DNA extraction, and quantification: ssDNA *vs*. dsDNA

Average DNA concentrations in treated samples and controls measured by the ds- and ssDNA Qubit assays are outlined in Table 1. Comparison of the ds- and ssDNA assay results, suggests that the HS dsDNA assay underestimates total DNA in the sample after treatment of NALC – NaOH decontaminated samples (Table 1). The dsDNA and ssDNA quantification assays both demonstrated that on average all treatment groups had a lower final DNA concentration following DNA extraction in comparison to the controls, and that both DNase and Benzonase treatments lead to a significant reduction in the total DNA concentration of each sample. This reduction suggests that there is significant amount of extracellular DNA in decontaminated sputum, and that the majority of this DNA is single stranded.

**Table 1:** Average DNA concentrations of sample groups as measure by Qubit ds DNA assay and Qubit ssDNA assay for the head-to-head comparison.

Significant differences between control and treatment groups were determined by unpaired t-test with Welch's correction. Three technical replicates were done per treatment group.
| Treatment | DNA concentration ng/ul | p value treatment vs control |
| --- | --- | --- |
| Qubit HS dsDNA assay |  |  |
| Control | 1,6 ± 0,32 | - |
| Wash | 1 ± 0,27 | 0,0491 |
| DNase | 0,37 ± 0,08 | 0,0146 |
| Benzonase | 0,38 ± 0,09 | 0,0151 |
| Qubit ssDNA assay |  |  |
| Control | 5,32 ± 1,52 | - |
| Wash | 2,7 ± 1,36 | 0,0911 |
| DNase | 0,55 ± 0,30 | 0,0297 |
| Benzonase | 0,51 ± 0,21 | 0,0283 |

### qPCR quantification of *Mtb* and host DNA

Analysing the quantitative PCR results for the control demonstrated a higher average copy number of the host genome (6 168 ± 1 638 copies/ng) relative to the *Mtb* target (212,3 ± 59,4 copies/ng), with an unfavourable ratio of 1:29 target versus host genomic copies. A simple wash of the sample or its treatment according to the DNase protocol (which includes 2 washes with DNAse buffer as well as the DNase treatment, Figure 1) led to a significant improvement in the copy numbers present for Mtb target DNA and a significant reduction in contaminant host DNA, as well as an improved ratio of Mtb DNA:host DNA when comparing the treatment conditions to the control (Table 2).

**Table 2:**
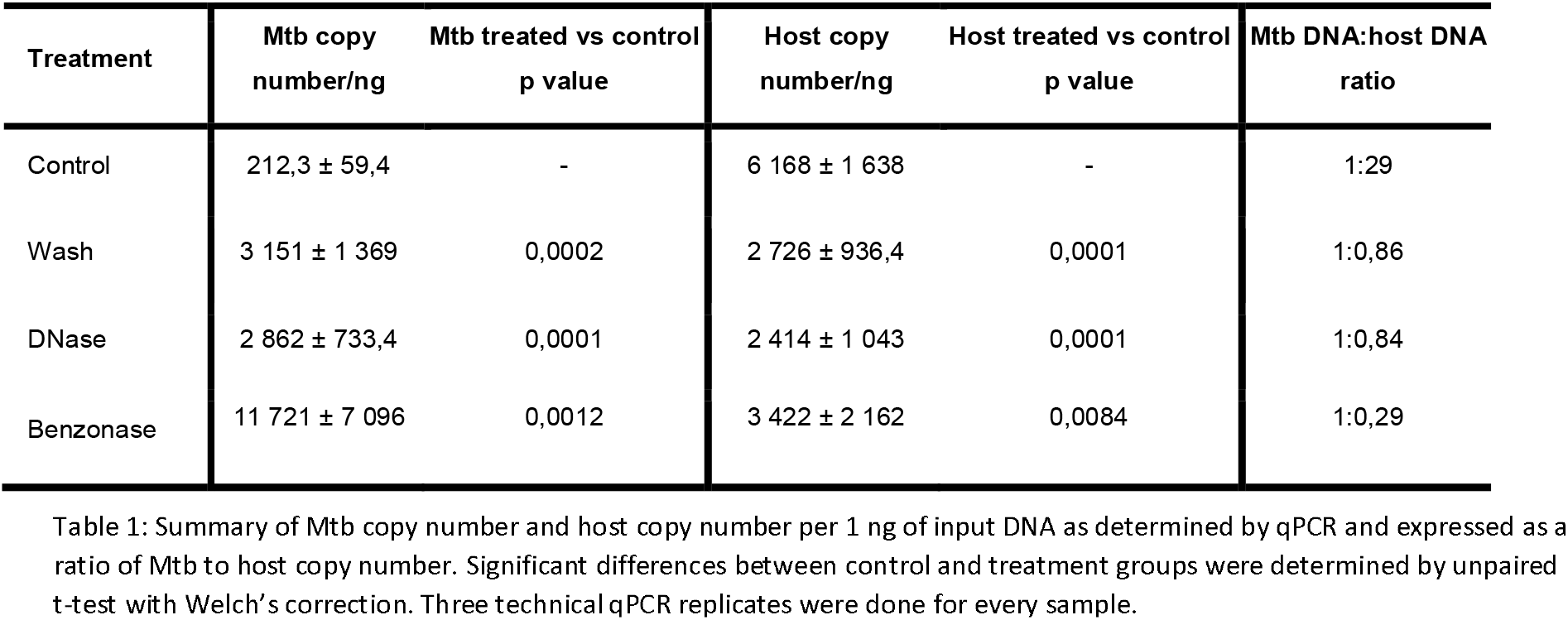
Decontamination by wash *vs*. DNase *vs*. benzonase prior to cell lysis: Copy number/ng of input DNA and ratios of Mtb:host are computed from qPCR performed on same starting material that was split four-fold between the control and three test protocols.

No significant difference was observed between either the Mtb copy number (p = 0,6) or the host copy number (p = 0,5), or the ratio of Mtb DNA:host DNA (1:0,86 vs 1:0,84) when directly comparing the Washed and DNase treated samples. This suggests that the impact is mainly due to the wash steps of the cellular material. Benzonase treatment showed enrichment for Mtb DNA, resulting in a 100-fold enrichment in the ratio of Mtb DNA:host DNA using benzonase (1:0.29) over the control (1:29) (Table 1). In summary, a simple wash of the decontaminated sample leads to an improved ratio of Mtb to host DNA, but additional pre-treatment with benzonase resulted in further enrichment of the Mtb target.

### Enrichment with TWIST target capture

#### Limit of detection

To initially evaluate the TWIST target capture system, we assessed the limit of detection (LoD) using a series of samples spiked with H37Rv DNA at progressively lower dilutions. The target capture and enrichment system successfully captured and detected Mtb down to a 100 genome copies. The proportion of the reference genome captured with the custom panel was 99,99% for the entire limit of detection range except for the sample equivalent to 10 genomes. We summarized the amount of data sequenced, raw read counts, read quality, insert size, coverage, duplication and error rates as well as the proportion of the reference genome covered at 5x and 10x (Supplementary Table 6). Coverage across the reference genome demonstrated high uniformity of capture (Supplementary Figure 4). Genomic windows where coverage deviated from the median (>1.5x) were visually inspected by IGV. The regions where an increase in coverage was observed were mainly attributed to insertion sequences (*IS6110, IS1081* and *IS1557*), while regions exhibiting an overall decrease in coverage were attributed to repeat regions (PE, PPE and Esx).

#### Direct sputum samples

DNA extracted from each of the treatment conditions for the wash versus DNase versus benzonase experiment was included in the TWIST target capture system. Classification of reads with kraken2 revealed the percentage of reads classifying as Mtb to be 89,28%, 0,42% from host and 10,3% from other host commensals (Figure 2). To assess the effectiveness of enrichment, the Benzonase treated sample was also sequenced directly because it had the highest enrichment ratio by qPCR. For the directly sequenced sample only 1,18% of reads were classified as target Mtb reads compared to 90,95% for the enriched sample. In addition to this 4,24 and 44,58% of reads were classified as belonging to the host or host commensals compared to 0,32 and 8,73% for the enriched sample respectively. The average depth of coverage obtained for the treated samples were 8x (Wash), 16x (DNase treated) and 22x (Benzonase treated) respectively, and the proportion of the genome covered at least 1x was 99.45 % on average with > 80% of the genome covered 5x (Supplementary Table 6). The total amount of data sequenced per sample ranged between 0,05 - 0,13 GB and the expected coverage per 1GB sequenced can be found in Supplementary Table 7.

**Figure 2:**
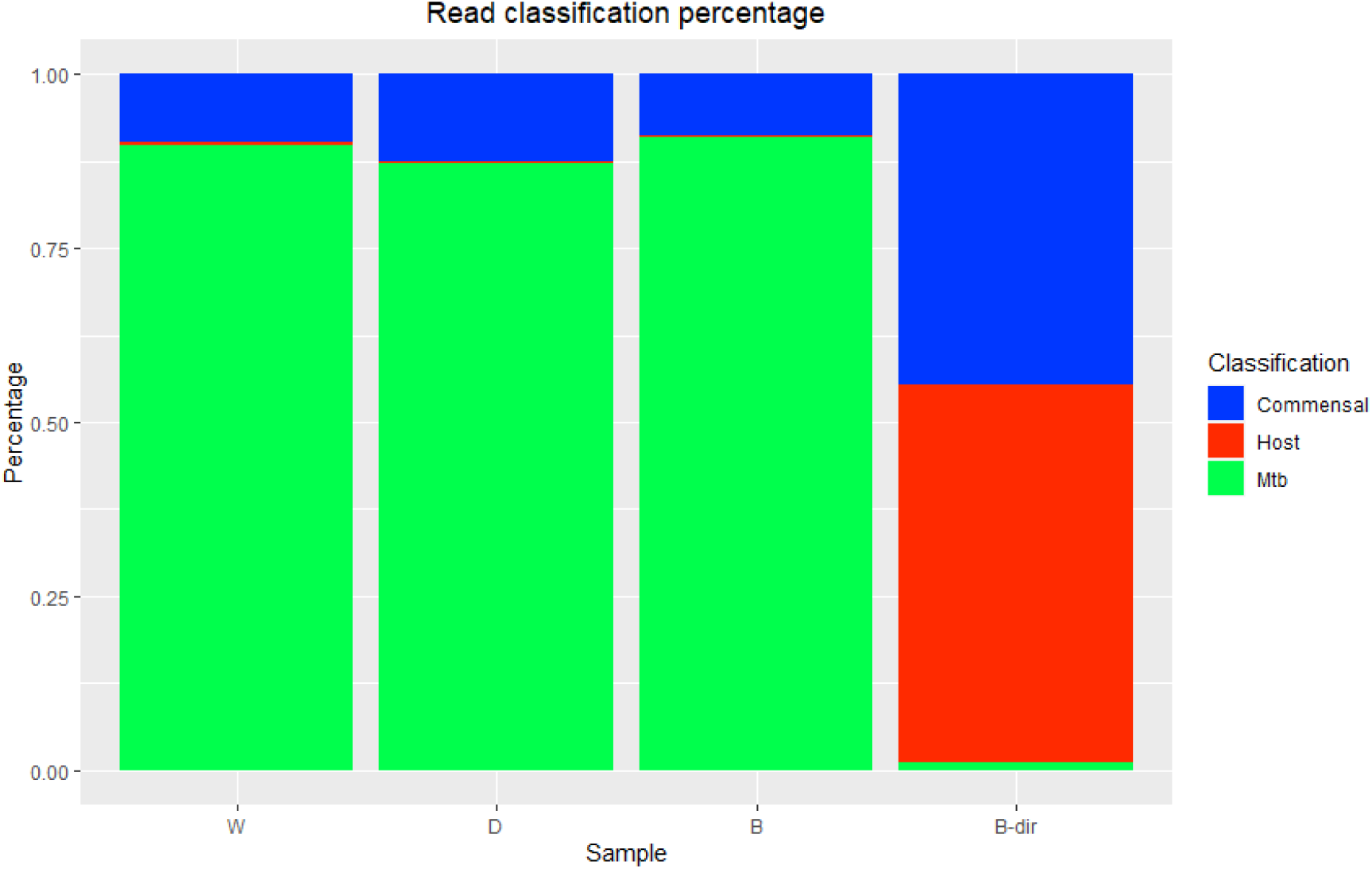
Percentage reads classified as either Mtb, host or other contaminating DNA. W (wash only sample 1 - enriched), D (DNase treated sample 1 - enriched), B (Benzonase treated sample 1 - enriched) and B-dir (Benzonase sample 1 - unenriched).

Using IGV (Robinson et al., 2011), we visually inspected Genomic regions with extreme coverage (>1.5x) in the benzonase treated sample (Supplementary Figure 5). Similar to the LoD experiments, an increase in coverage was observed at insertion sequences (*IS6110, IS1081* and *IS1557*), while repeat regions (*PE*, *PPE* and *esX*) exhibited a decrease in overall coverage, for example *IS6610* elements had an average coverage of 77x and as a representative example *PE_PGRS4* had an average coverage of 3.8x. There was also a significant increase in the average coverage (891x) observed across the *rrs* and *rrl* genes. Several other regions conserved among bacterial species such as *clpB, HSP, rpoB, rpoC, tuf, rpsc* and *aspT* were also found to have more than expected coverage ranging between 5 and to 9-fold higher than the genomic average.

Given the high conservation of the *rrs/rrl* regions and the aforementioned genes, we hypothesized that the observed high coverage is due to off target capture of these elements from non-Mtb respiratory bacteria contaminating the sample. We classified all bacterial reads mapping to the *rrs-rrl* region (13,50 % of total sequencing reads) using the Kraken2 (Lu et al., 2022). Only 4,83 % of these reads were classified as Mtb confirming that the extreme coverage was due to off target capture. The six most abundant source genera for the rrsl/rrl reads overrepresented respiratory flora including Streptococcus (19,68%), Streptomyces (5,62%), Arthrobacter (3,92%), Rhodococcus (1,42%), Bfidiobacterium (1,52%) and Staphylococcus (1,63%). We then used Kraken2 to remove contaminating DNA across the whole genome, by excluding all reads not classified as Mtb *complex* (9,05 % of total) prior to mapping to the H37Rv reference genome. This corrected the non-uniformity of coverage in all regions except for *16S (rrs) and 23S (rrl)*, that were still covered at 2,5-fold across both genes and up to 15-fold the genomic average in certain regions.

## Discussion

Sputum specimens collected for Mtb detection remain challenging for DSS due to the large quantities of contaminating host DNA they contain. We aimed to evaluate two approaches, both separately and in conjunction, to facilitate DSS. Results reveal that host DNA can be successfully depleted by washing the samples or treating the samples with DNase or benzonase, leading to a significantly improved ratio of Mtb DNA:host DNA. Ideal results the required target capture and enrichment as well, which we included via a DNA probe based capture system from Twist Bioscience (Brown et al., 2015; Doyle et al., 2018; Goig et al., 2020; Nimmo et al., 2019). Finally, we also report that dsDNA assays do not accurately estimate DNA concentrations when working with NaOH decontaminated sediments, underestimating DNA concentration compared to a ssDNA assay.

In standard procedures, sputum samples are decontaminated under alkaline conditions (pH 12-14) prior to downstream processing for Mtb culture (Votintseva *et al*., 2015). Alkaline conditions (pH > 9) such as those generated by sodium hydroxide, denature DNA leading to large amounts of ssDNA in the sample. This observation is important since techniques like qPCR and NGS rely on accurate DNA quantification in the input sample and incorrect quantification could also impact the final results (Wang *et al*., 2014; He *et al*., 2018). Alkaline conditions and heat inactivation, as used in our protocol, can both lyse contaminating host cells and gram-negative organisms. This then causes the majority of the contaminating DNA to become extracellular and available for enzymatic degradation (Hasan *et al*., 2016; Shehadul Islam *et al*., 2017), meaning washing the sediment sample prior to DNA extraction will remove most of these contaminants. This is congruent with previous studies where the application of a saline wash or treatment with the MolYsis kit (Molzym, Germany), reduced contaminating DNA (Votintseva *et al*., 2015; Votintseva *et al*., 2017).

The benzonase treatment protocol demonstrated a significant enrichment effect. We initially hypothesized that benzonase was superior to a simple wash or DNase treatment because it more effectively removes extracellular contaminating ssDNA in addition to dsDNA, RNA and DNA:RNA hybrids, while DNase targets only dsDNA (Sutton *et al*., 1997; Liu *et al*., 2019; Amar *et al*., 2021). However, there was only a small difference in total DNA by the Qubit ssDNA between the DNase and benzonase treated samples (0,55 ± 0,302 ng/μl and 0,51 ± 0,213 ng/μl respectively), arguing that more effective ssDNA degradation is not the only reason for the benzonase effect. An alternative explanation is that the benzonase treatment protocol results in more effective Mtb lysis than the other two protocols, thus enhancing recovery of Mtb. The exact mechanisms behind the observed enrichment effect will be the focus of future studies.

Although a positive enrichment effect was observed, target capture and enrichment was still necessary because of the low levels of target DNA from DSS. (Votintseva *et al*., 2017; Nimmo *et al*., 2019; Goig *et al*., 2020a). Our results demonstrate that the Twist kit can handle low input amounts of DNA down to 100 Mtb genome copies (0,45 pg of Mtb input DNA), making it an important breakthrough necessary for success for DSS. After Twist enrichment, WGS quality, expected coverage and uniformity was high. Sequencing data demonstrated a very low proportion of duplicate reads (median 0,43%) in relation to a previous study which reported an extremely high proportion of duplicates (median 80%) for samples enriched with RNA baits (Goig et al., 2020a). In addition to the low propensity to capture host DNA, we have demonstrated the effectiveness of the Twist target capture and enrichment system to enrich for the Mtb target from 1,18 % in the directly sequenced sample to 90,95% in the enriched. We also found no added benefit with additional pre-treatments prior to use of the Twist kit other than recommended wash steps as the proportion of reads attributed ranged between 87,12 - 90,95 % and were sufficient for the recovery of the target genome for all 3 treatment conditions. We did identify off-target capture (9,05% of total reads) for highly conserved domains, especially the *rRNA* regions from contaminating respiratory flora, but not from host DNA. The off-target capture of rRNA elements is not unique to the evaluated enrichment platform and several previous studies utilising RNA baits have also reported off-target capture and poor uniformity in these regions (Brown *et al*., 2015; Nimmo *et al*., 2019).

### Limitations

The current study has limitations. The limit of detection experiments consisted of pure H37Rv in culture media and not in sputum and hence excluded the effects of contaminating DNA. This allowed us to isolate the effect of the inoculum and study sequencing uniformity independent of contamination. For the sputum based experiments, we used pooled sputum sediments that increased the risk of contamination but was needed to generate enough sample volume for protocol comparisons and technical replicates (Burdz *et al*., 2003; Asmar & Drancourt, 2015). Due to the limited amount of DNA that can be extracted from this sample type we opted to only quantify host and target DNA, thus omitting the quantification of overall bacterial load. This decision was made to prioritise host DNA quantification as Host DNA is generally the primary contributing contaminant in clinical specimens ((Brown et al., 2015; Doyle et al., 2018; Nimmo et al., 2019; Goig et al., 2020).

We note that our input samples were generated from a pool of several sediments. There was thus an expected higher risk of contamination and constituted a more extreme test of the specificity of capture compared with a single patient isolate. Off-target capture was readily addressable bioinformatically by excluding non-Mtbc reads. Prior work on DSS also suggested removing or masking the rRNA genes in WGS data prior to variant calling (Brown *et al*., 2015; Nimmo *et al*., 2019; George *et al*., 2020). Despite the bioinformatics solution, off target capture increases cost and decreases efficiency of DSS. In the future, redesign of the panel to exclude or de-enrich these regions may be considered. The probe panel was supplemented by 4-fold coverage for homoplastic SNVs in drug resistance regions, 91 of which are found in the rrs and rrl genes, potentially exacerbating the extent of off target capture. Alternatively, refinement of current protocols to prevent nonspecific binding e.g. by capture at higher temperatures, may also reduce off-target capture (Paijmans et al., 2016).

### Conclusions

We demonstrate that pre-processing with a benzonase enrichment protocol to remove contaminating extracellular DNA prior to cell lysis and DNA extraction has a positive effect for the enrichment of *Mtb* DNA in decontaminated sediments, but the exact mechanisms by which enrichment is facilitated is still unknown. In addition, we demonstrate the utility and low limit of detection of the DNA probe-based Twist target capture and enrichment system for enrichment and sequencing of Mtb from clinical sputum sediments.

## Supporting information

Supplemental folder - 1

