## Supplementary file -1 for "Assessment and validation of enrichment and target capture approaches to improve *Mycobacterium tuberculosis* WGS from direct patient samples"

**Supplementary appendix**

### Supplementary methods and results

#### Supplementary methods 1 – Enzymatic activity testing for DNase and Benzonase

DNase activity Benzonase activity was evaluated as depicted in supplementary Figure 2. Two pools of heat inactivated sediment were generated from routinely discarded TB positive sediments (NALC - NaOH decontaminated sputum samples – referred to as sediments). Each pool was split into several aliquots, which were then divided into groups of 3 replicates each per treatment condition. For the DNase evaluation, groups consisted of a treated, wash and control group, and for the Benzonase evaluation groups consisted of a treated and control group. Treatment for each group was done as described in the main text (Figure 1). Following respective treatments DNA was extracted and qPCR was performed as described under the Materials and Methods section.

#### Supplementary methods 2 – Validation of limit of detection dilution series

Considering the lower limit DNA concentrations used for the LoD were well below the quantification limit of the Qubit dsDNA HS Assay kit, qPCR was used to determine the accuracy of the dilution series generated. The starting sample as measured by Qubit was 4.52ng/µl or ~10^6^ H37Rv genomes and diluted 10-fold from there for each sample. Briefly 10 µl of starting sample was transferred to 90 µl of nuclease free H_2_O, pulse vortexed for 5 s, centrifuged and then incubated on ice for 5 min before the transfer to the next dilution in the series. This was repeated until the desired concentrations were reached for input into the Twist system as seen in Supplementary Table 5. qPCR was then preformed on the dilution series and observed for a linear reduction indicative of accurate dilution. qPCR was performed on the Quant Studio 5 (Thermofisher Scientific, USA), and the reaction set up was done as described under the Materials and Methods section, with Mtb specific primers and 4 µl of input material from each dilution.

#### Supplementary methods 3 – Estimation of costs

Current estimates on prices for target capture and enrichment were calculated as of January 2023. Pricing per reaction was calculated based on cost per reaction for library preparation and hybridisation. Estimates for the Roche and Agilent platforms were provided by local suppliers, and for the methods applied for Arbor biosciences’s capture and enrichment platform we relied on previous published works by Galo et al. 2020a. Calculations do not take into account: DNA extraction, DNA purification, prior Mtb quantification, personnel cost, quality control during library preparation or sequencing.

#### Supplementary results 1 – Enzymatic activity testing for DNase and Benzonase

Average DNA concentrations obtained for initial evaluations of DNase and Benzonase activity on NALC – NaOH decontaminated samples demonstrated that on average all treatment groups had a lower final DNA concentration following DNA extraction in comparison to the controls, and that both DNase and Benzonase treatments lead to a significant reduction in the total DNA concentration of each sample (Supplementary table 1 & 2). The same observation was also made when examining the results from the ssDNA assay kit. The reduction in overall yield of DNA when comparing enzymatically treated samples to control and wash sample groups indicate that a wash already has an effect in removing extracellular DNA, but that the addition of an enzyme lead to an even further reduction in the final yield of DNA. Analysing the quantitative PCR results for the initial pilot experiments demonstrated that just washing the sample prior to DNA extraction increased the ratio of Mtb to host DNA, but the addition of a DNase or benzonase treatment step resulted in an even higher ratio (Supplementary Table 3 and 4).

#### Supplementary results 2 – Validation of limit of detection dilution series

A linear decrease was observed following qPCR for the prepared LoD, indicative of accurate dilution (Supplementary Figure 1).

### Supplementary Tables

**Supplementary Table 1: Average DNA concentrations of sample groups as measure by Qubit ds DNA assay and Qubit ssDNA assay for the DNase activity test**

| **Treatment** | **DNA concentration ng/ul** | **p value treatment vs control** |
| --- | --- | --- |
|  | **Qubit HS dsDNA assay** | |
| Control | 5,05 ± 0,41 | - |
| Wash | 2,92 ± 0,2 | 0,0045 |
| DNase | 0,5 ± 0,14 | 0,0012 |
|  | **Qubit ssDNA assay** | |
| Control | 19,73 ± 0,59 | - |
| Wash | 10,47 ± 1,1 | 0,0009 |
| DNase | 0,56 ± 0,12 | 0,0002 |

**Supplementary Table 2: Average DNA concentrations of sample groups as measure by Qubit ds DNA assay and Qubit ssDNA assay for the Benzonase activity test**

| **Treatment** | **DNA concentration ng/ul** | **p value treatment vs control** |
| --- | --- | --- |
|  | **Qubit HS dsDNA assay** | |
| Control | 6,64 ± 0,31 | - |
| DNase | 0,21 ± 0,05 | 0,0001 |
|  | **Qubit ssDNA assay** | |
| Control | 23,4 ± 2,69 | - |
| DNase | 0,19 ± 0,036 | 0,0044 |

**Supplementary Table 3: Copy number and ratio comparison for the DNase optimisation experiment**

Supplementary Table 3: Summary of *Mtb* copy number and host copy number per ng of input DNA as determined by qPCR and expressed as a ratio of *Mtb* to host copy number. Significant differences between control and treatment groups were determined by unpaired t-test with Welch’s correction. 3 technical qPCR replicates were done for every sample.

| **Treatment** | ***Mtb* copy number** | ***Mtb* treated vs control p value** | **Host copy number** | **Host treatment vs control p value** | ***Mtb* DNA:host DNA ratio** |
| --- | --- | --- | --- | --- | --- |
| Control | 1 713 ± 624 | - | 10 424 ± 1 521 | - | 1:5,34 |
| Wash | 2 474 ± 556,3 | 0,0149 | 5 539 ± 1561 | 0,0001 | 1:2,25 |
| DNase | 1 950 ± 312 | 0,3284 | 84,45 ± 18,97 | 0,0001 | 1:0,05 |

**Supplementary Table 4: Copy number and ratio comparison for the Benzonase optimisation experiment**

Supplementary Table 4: Summary of *Mtb* copy number and host copy number per ng of input DNA as determined by qPCR and expressed as a ratio of *Mtb* to host copy number. Significant differences between control and treatment groups were determined by unpaired t-test with Welch’s correction. 2 technical qPCR replicates were done for every sample.

| **Treatment** | ***Mtb* copy number** | ***Mtb* treated vs control p value** | **Host copy number** | **Host treatment vs control p value** | ***Mtb* DNA:host DNA ratio** |
| --- | --- | --- | --- | --- | --- |
| Control | 106 ± 23,11 | - | 1 867 ± 396,4 | - | 1:5,34 |
| Benzonase | 3028 ± 1752 | 0,3284 | 1 560 ± 960,6 | 0,3284 | 1:0,05 |

**Supplementary Table 5: Summary of samples for sequencing using the Twist target capture and enrichment system**

| **Sample** | **Input into library prep (ng)** | **Mtb input into library prep (ng)** | **Genomes** |
| --- | --- | --- | --- |
| H37Rv - 10^5^ | 0,452 | 0,452 | 100 000 |
| H37Rv - 10^4^ | 0,0 452 | 0,0 452 | 10 000 |
| H37Rv - 10^3^ | 0,00 452 | 0,00 452 | 1 000 |
| H37Rv - 10^2^ | 0,000 452 | 0,000 452 | 100 |
| H37Rv - 10 | 0,0 000 452 | 0,0 000 452 | 10 |
| W | 4,5 | 0,033 | 7298,73 |
| D | 4,54 | 0,044 | 9748,71 |
| B | 4,28 | 0,432 | 95701,34 |

**Supplementary Table 6: Quality control and coverage estimates**

| **Sample** | **Library prep** | **Data sequenced (Gb)** | **Raw read count** | **% Reads > Q30** | **Reads mapped to reference** | **Average insert size** | **Average depth of coverage** | **Duplication %** | **Error rate %** | **Proportion of reference genome recovered at 1x (%)** | **- 5x** | **- 10x** |
| --- | --- | --- | --- | --- | --- | --- | --- | --- | --- | --- | --- | --- |
| H37Rv - 10^5^ | Twist | 4,54 | 30 445 936 | 88,8 | 29 081 846 | 182 | 873±193 | 3,37 | 0,55 | 99,99 | 99,99 | 99,99 |
| H37Rv - 10^4^ | Twist | 3,87 | 25 996 020 | 88,3 | 24 861 691 | 187 | 749±160 | 3,13 | 0,58 | 99,99 | 99,99 | 99,99 |
| H37Rv - 10^3^ | Twist | 2,29 | 15 388 576 | 88,4 | 14 812 085 | 190 | 433±93 | 6,15 | 0,56 | 99,99 | 99,99 | 99,99 |
| H37Rv - 10^2^ | Twist | 0,33 | 2 232 812 | 88,2 | 2 151 371 | 187 | 62±17 | 6,4 | 0,57 | 99,99 | 99,97 | 99,91 |
| H37Rv - 10 | Twist | - | 218 | - | - | - | - | - | - | - | - | - |
| W | Twist | 0,05 | 309 150 | 88,8 | 268 127 | 175 | 8±4 | 0,42 | 0,62 | 98,63 | 80,62 | 34,13 |
| D | Twist | 0,09 | 636 268 | 89,1 | 534 797 | 168 | 16±7 | 0,41 | 0,6 | 99,79 | 97,72 | 85,27 |
| B | Twist | 0,13 | 843 304 | 88,7 | 746 841 | 166 | 22±9 | 0,45 | 0,65 | 99,92 | 99,35 | 95,87 |
| B-dir | Illumina | 2,46 | 16 369 512 | 90,8 | 191 475 | 139 | 5±3 | 0,24 | 0,58 | 95.68 | 52.93 | 8.56 |

**Supplementary Table 7: Expected coverage per 1GB of data sequenced**

| **Sample** | **GB total** | **Coverage (Median)** | **Expected coverage per 1 GB output** |
| --- | --- | --- | --- |
| **W** | 0,05 | 8 | 173,67 |
| **D** | 0,09 | 16 | 168,77 |
| **B** | 0,13 | 22 | 175,09 |
| **B-dir** | 2,46 | 5 | 2,03 |

### Supplementary Figures

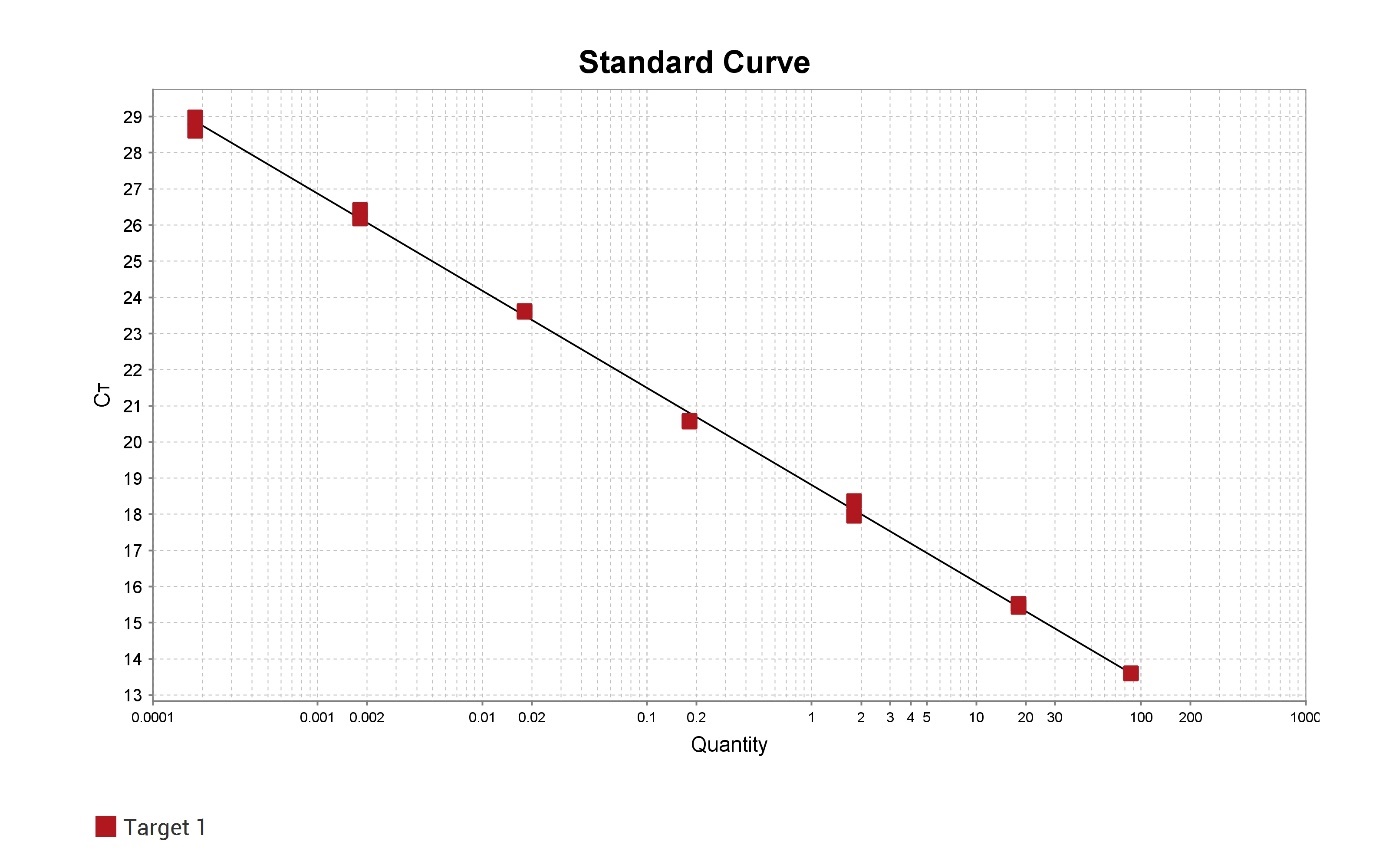

Supplementary Figure 1: Standard curve generated to validate dilution accuracy of the generated H37Rv limit of detection.

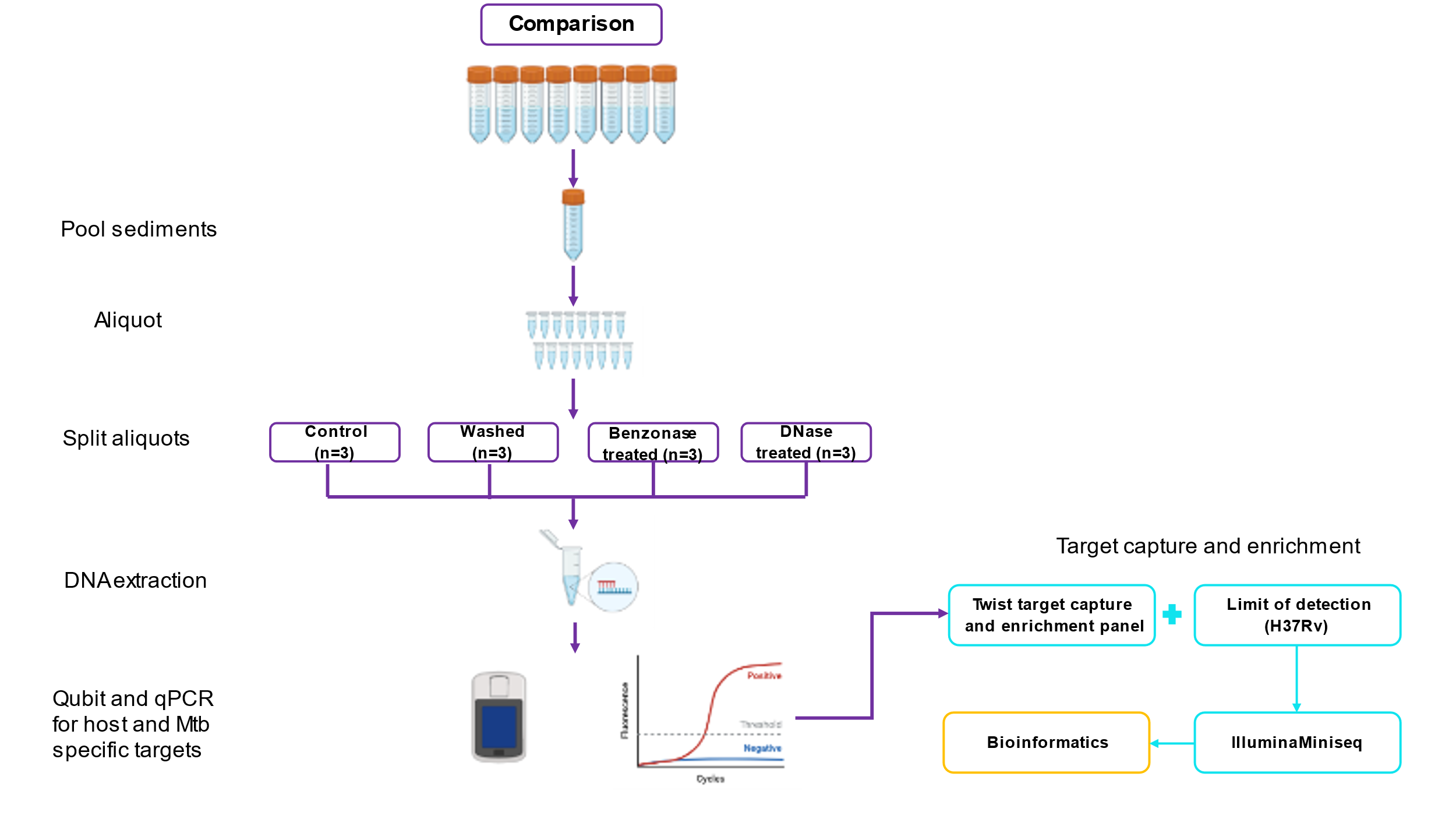

Supplementary Figure 2: Workflow starting from enzymatic depletion of contaminating DNA to sequencing with the Twist target capture and enrichment system for the H37Rv limit of detection as well as selected samples following enzymatic depletion of contaminating DNA.

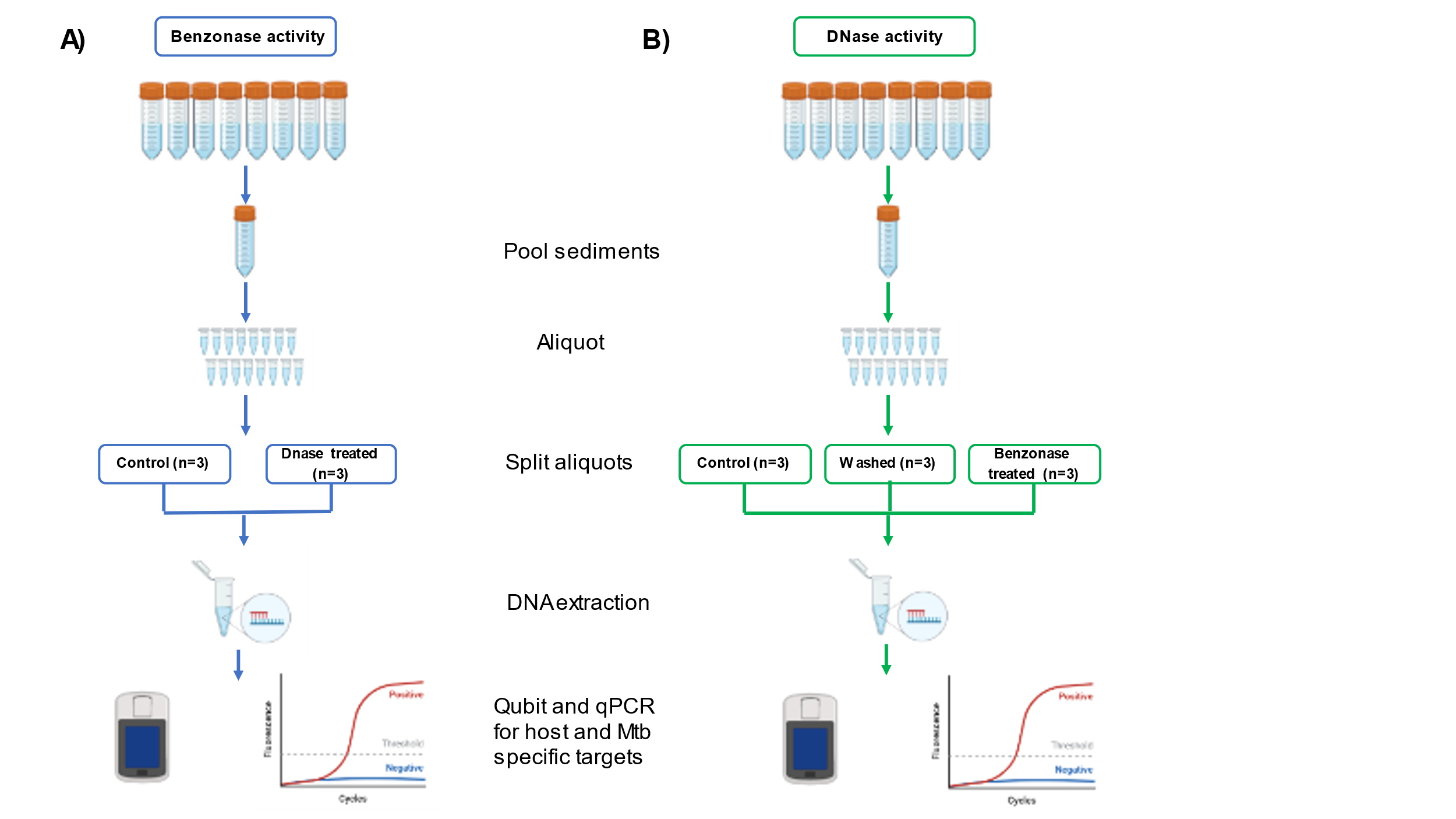

Supplementary Figure 3: Workflow for initial enzymatic activity estimation for DNase and Benzonase treatment workflows.

Coverage across reference - H37Rv - 105

|  | Normalised |
| --- | --- |
| 2206.5 | 1.28272 |
| 6618.5 | 1.55596 |
| 11030.5 | 1.17347 |
| 15442.5 | 1.01347 |
| 19854.5 | 1.0201 |
| 24266.5 | 0.995021 |
| 28678.5 | 1.02998 |
| 33090.5 | 1.04622 |
| 37502.5 | 1.04152 |
| 41914.5 | 1.07204 |
| 46326.5 | 1.05323 |
| 50738.5 | 1.06637 |
| 55150.5 | 1.10872 |
| 59562.5 | 1.13259 |
| 63974.5 | 1.23753 |
| 68386.5 | 1.05194 |
| 72798.5 | 1.10756 |
| 77210.5 | 1.00518 |
| 81622.5 | 1.07721 |
| 86034.5 | 1.03768 |
| 90446.5 | 0.930817 |
| 94858.5 | 0.970306 |
| 99270.5 | 0.918561 |
| 103682 | 1.1047 |
| 108094 | 1.18349 |
| 112506 | 1.13356 |
| 116918 | 1.06105 |
| 121330 | 1.01787 |
| 125742 | 1.00186 |
| 130154 | 0.969651 |
| 134566 | 1.0642 |
| 138978 | 1.16896 |
| 143390 | 1.06126 |
| 147802 | 1.05441 |
| 152214 | 0.982323 |
| 156626 | 1.05342 |
| 161038 | 1.02663 |
| 165450 | 0.967611 |
| 169862 | 0.94688 |
| 174274 | 1.04018 |
| 178686 | 0.97601 |
| 183098 | 0.997143 |
| 187510 | 0.974552 |
| 191922 | 1.02858 |
| 196334 | 1.1297 |
| 200746 | 0.996302 |
| 205158 | 1.07189 |
| 209570 | 1.07833 |
| 213982 | 1.04769 |
| 218394 | 1.0649 |
| 222806 | 1.08005 |
| 227218 | 1.03385 |
| 231630 | 0.995195 |
| 236042 | 0.925261 |
| 240454 | 0.950748 |
| 244866 | 0.933273 |
| 249278 | 1.01504 |
| 253690 | 1.0201 |
| 258102 | 1.05527 |
| 262514 | 0.943078 |
| 266926 | 0.956406 |
| 271338 | 1.01286 |
| 275750 | 0.940256 |
| 280162 | 0.949723 |
| 284574 | 0.909037 |
| 288986 | 0.960478 |
| 293398 | 0.992323 |
| 297810 | 1.16418 |
| 302222 | 1.123 |
| 306634 | 1.0014 |
| 311046 | 0.965651 |
| 315458 | 0.953794 |
| 319870 | 0.902749 |
| 324282 | 0.925361 |
| 328694 | 0.990106 |
| 333106 | 0.875592 |
| 337518 | 0.600138 |
| 341930 | 0.887132 |
| 346342 | 0.947909 |
| 350754 | 0.94377 |
| 355166 | 0.841466 |
| 359578 | 0.950389 |
| 363990 | 0.917266 |
| 368402 | 0.975426 |
| 372814 | 0.996661 |
| 377226 | 1.07079 |
| 381638 | 1.12456 |
| 386050 | 1.0768 |
| 390462 | 1.07763 |
| 394874 | 1.02818 |
| 399286 | 1.07005 |
| 403698 | 1.04164 |
| 408110 | 0.992455 |
| 412522 | 1.10562 |
| 416934 | 1.08412 |
| 421346 | 1.07283 |
| 425758 | 0.901513 |
| 430170 | 1.03558 |
| 434582 | 0.915756 |
| 438994 | 0.989312 |
| 443406 | 0.90763 |
| 447818 | 1.00336 |
| 452230 | 0.954895 |
| 456642 | 0.844747 |
| 461054 | 0.908629 |
| 465466 | 0.857451 |
| 469878 | 0.930289 |
| 474290 | 0.958269 |
| 478702 | 1.01331 |
| 483114 | 1.17593 |
| 487526 | 1.1821 |
| 491938 | 1.03482 |
| 496350 | 0.944353 |
| 500762 | 0.93107 |
| 505174 | 1.03614 |
| 509586 | 0.927301 |
| 513998 | 0.897871 |
| 518410 | 0.954122 |
| 522822 | 0.913046 |
| 527234 | 0.962703 |
| 531646 | 0.887678 |
| 536058 | 0.980126 |
| 540470 | 1.02796 |
| 544882 | 1.02772 |
| 549294 | 1.03714 |
| 553706 | 1.07232 |
| 558118 | 1.08087 |
| 562530 | 1.09618 |
| 566942 | 0.948033 |
| 571354 | 0.887683 |
| 575766 | 0.933406 |
| 580178 | 0.998218 |
| 584590 | 0.897754 |
| 589002 | 0.982222 |
| 593414 | 1.0334 |
| 597826 | 1.1018 |
| 602238 | 1.07423 |
| 606650 | 1.07493 |
| 611062 | 0.958666 |
| 615474 | 0.875833 |
| 619886 | 0.951293 |
| 624298 | 0.76302 |
| 628710 | 0.81272 |
| 633122 | 0.845973 |
| 637534 | 0.948324 |
| 641946 | 0.991841 |
| 646358 | 0.986584 |
| 650770 | 1.10283 |
| 655182 | 1.09505 |
| 659594 | 1.01337 |
| 664006 | 1.02248 |
| 668418 | 0.925493 |
| 672830 | 0.710319 |
| 677242 | 0.940374 |
| 681654 | 1.06755 |
| 686066 | 1.1472 |
| 690478 | 1.04886 |
| 694890 | 1.07179 |
| 699302 | 0.914371 |
| 703714 | 0.9366 |
| 708126 | 0.920199 |
| 712538 | 0.838825 |
| 716950 | 0.912547 |
| 721362 | 0.945527 |
| 725774 | 0.861236 |
| 730186 | 0.976329 |
| 734598 | 0.975302 |
| 739010 | 1.06222 |
| 743422 | 0.868344 |
| 747834 | 0.738873 |
| 752246 | 0.94925 |
| 756658 | 1.01764 |
| 761070 | 1.27627 |
| 765482 | 1.22244 |
| 769894 | 1.00464 |
| 774306 | 1.00579 |
| 778718 | 1.06968 |
| 783130 | 1.12292 |
| 787542 | 1.00974 |
| 791954 | 1.00829 |
| 796366 | 1.02741 |
| 800778 | 1.12103 |
| 805190 | 0.958071 |
| 809602 | 1.00259 |
| 814014 | 0.96624 |
| 818426 | 0.981326 |
| 822838 | 0.878793 |
| 827250 | 1.01778 |
| 831662 | 0.846702 |
| 836074 | 0.770055 |
| 840486 | 0.853944 |
| 844898 | 0.874802 |
| 849310 | 0.936318 |
| 853722 | 1.03159 |
| 858134 | 1.00592 |
| 862546 | 0.937554 |
| 866958 | 0.946494 |
| 871370 | 0.885219 |
| 875782 | 1.10463 |
| 880194 | 1.0733 |

Coverage across reference - H37Rv - 104

|  | Norm |
| --- | --- |
| 2206.5 | 1.25157 |
| 6618.5 | 1.5628 |
| 11030.5 | 1.16599 |
| 15442.5 | 0.994616 |
| 19854.5 | 0.997944 |
| 24266.5 | 0.993339 |
| 28678.5 | 1.02653 |
| 33090.5 | 1.02226 |
| 37502.5 | 1.02961 |
| 41914.5 | 1.09897 |
| 46326.5 | 1.06607 |
| 50738.5 | 1.0635 |
| 55150.5 | 1.09771 |
| 59562.5 | 1.10934 |
| 63974.5 | 1.22496 |
| 68386.5 | 1.04415 |
| 72798.5 | 1.12389 |
| 77210.5 | 1.01464 |
| 81622.5 | 1.08796 |
| 86034.5 | 1.04663 |
| 90446.5 | 0.941247 |
| 94858.5 | 0.986603 |
| 99270.5 | 0.932033 |
| 103682 | 1.13953 |
| 108094 | 1.16046 |
| 112506 | 1.11014 |
| 116918 | 1.05739 |
| 121330 | 1.0227 |
| 125742 | 1.02276 |
| 130154 | 0.991854 |
| 134566 | 1.06067 |
| 138978 | 1.17739 |
| 143390 | 1.06173 |
| 147802 | 1.05783 |
| 152214 | 0.97551 |
| 156626 | 1.04632 |
| 161038 | 1.03357 |
| 165450 | 0.961532 |
| 169862 | 0.942518 |
| 174274 | 1.05718 |
| 178686 | 0.988896 |
| 183098 | 1.0004 |
| 187510 | 0.955719 |
| 191922 | 1.03335 |
| 196334 | 1.1119 |
| 200746 | 0.980702 |
| 205158 | 1.04447 |
| 209570 | 1.07455 |
| 213982 | 1.05006 |
| 218394 | 1.06368 |
| 222806 | 1.07485 |
| 227218 | 1.03338 |
| 231630 | 0.993876 |
| 236042 | 0.907601 |
| 240454 | 0.952544 |
| 244866 | 0.917751 |
| 249278 | 1.01439 |
| 253690 | 1.02564 |
| 258102 | 1.05214 |
| 262514 | 0.936252 |
| 266926 | 0.976824 |
| 271338 | 1.00168 |
| 275750 | 0.956765 |
| 280162 | 0.920944 |
| 284574 | 0.898187 |
| 288986 | 0.949389 |
| 293398 | 0.955322 |
| 297810 | 1.14583 |
| 302222 | 1.12573 |
| 306634 | 0.99014 |
| 311046 | 0.96433 |
| 315458 | 0.964347 |
| 319870 | 0.913756 |
| 324282 | 0.919427 |
| 328694 | 0.989949 |
| 333106 | 0.902304 |
| 337518 | 0.640672 |
| 341930 | 0.882654 |
| 346342 | 0.960784 |
| 350754 | 0.944844 |
| 355166 | 0.852098 |
| 359578 | 0.944232 |
| 363990 | 0.925341 |
| 368402 | 0.952659 |
| 372814 | 0.962517 |
| 377226 | 1.06957 |
| 381638 | 1.14467 |
| 386050 | 1.05178 |
| 390462 | 1.08959 |
| 394874 | 1.0328 |
| 399286 | 1.06052 |
| 403698 | 1.01997 |
| 408110 | 1.00691 |
| 412522 | 1.10778 |
| 416934 | 1.10423 |
| 421346 | 1.05787 |
| 425758 | 0.883928 |
| 430170 | 1.00714 |
| 434582 | 0.891156 |
| 438994 | 0.972774 |
| 443406 | 0.902293 |
| 447818 | 1.00625 |
| 452230 | 0.956206 |
| 456642 | 0.834723 |
| 461054 | 0.910028 |
| 465466 | 0.883746 |
| 469878 | 0.919135 |
| 474290 | 0.959105 |
| 478702 | 1.00866 |
| 483114 | 1.16895 |
| 487526 | 1.17249 |
| 491938 | 1.03373 |
| 496350 | 0.943217 |
| 500762 | 0.94171 |
| 505174 | 1.06065 |
| 509586 | 0.918344 |
| 513998 | 0.927519 |
| 518410 | 0.953638 |
| 522822 | 0.921562 |
| 527234 | 0.962154 |
| 531646 | 0.890269 |
| 536058 | 0.971765 |
| 540470 | 1.01966 |
| 544882 | 1.05098 |
| 549294 | 1.04793 |
| 553706 | 1.05629 |
| 558118 | 1.0545 |
| 562530 | 1.06878 |
| 566942 | 0.957795 |
| 571354 | 0.88877 |
| 575766 | 0.93803 |
| 580178 | 1.00548 |
| 584590 | 0.912298 |
| 589002 | 0.985769 |
| 593414 | 1.03394 |
| 597826 | 1.11554 |
| 602238 | 1.0879 |
| 606650 | 1.08276 |
| 611062 | 0.963487 |
| 615474 | 0.89655 |
| 619886 | 0.932941 |
| 624298 | 0.776307 |
| 628710 | 0.860034 |
| 633122 | 0.856035 |
| 637534 | 0.957371 |
| 641946 | 0.969318 |
| 646358 | 0.998444 |
| 650770 | 1.13825 |
| 655182 | 1.10624 |
| 659594 | 0.99824 |
| 664006 | 1.05667 |
| 668418 | 0.910539 |
| 672830 | 0.733731 |
| 677242 | 0.933392 |
| 681654 | 1.09803 |
| 686066 | 1.13254 |
| 690478 | 1.04091 |
| 694890 | 1.05631 |
| 699302 | 0.925578 |
| 703714 | 0.936562 |
| 708126 | 0.932332 |
| 712538 | 0.845877 |
| 716950 | 0.910325 |
| 721362 | 0.936321 |
| 725774 | 0.859674 |
| 730186 | 0.967902 |
| 734598 | 0.962287 |
| 739010 | 1.03877 |
| 743422 | 0.882177 |
| 747834 | 0.764889 |
| 752246 | 0.960685 |
| 756658 | 1.02516 |
| 761070 | 1.26168 |
| 765482 | 1.19742 |
| 769894 | 1.00845 |
| 774306 | 1.00396 |
| 778718 | 1.03522 |
| 783130 | 1.09621 |
| 787542 | 1.02637 |
| 791954 | 1.021 |
| 796366 | 1.02105 |
| 800778 | 1.11522 |
| 805190 | 0.949181 |
| 809602 | 0.981133 |
| 814014 | 0.964311 |
| 818426 | 0.979117 |
| 822838 | 0.878933 |
| 827250 | 0.995107 |
| 831662 | 0.856236 |
| 836074 | 0.786873 |
| 840486 | 0.872666 |
| 844898 | 0.888429 |
| 849310 | 0.918617 |
| 853722 | 1.01221 |
| 858134 | 0.989888 |
| 862546 | 0.956304 |
| 866958 | 0.954088 |
| 871370 | 0.873884 |
| 875782 | 1.09158 |
| 880194 | 1.0719 |

Normalised sequencing depth

Coverage across reference - H37Rv - 103

|  | Norm |
| --- | --- |
| 2206.5 | 1.22324 |
| 6618.5 | 1.55997 |
| 11030.5 | 1.15615 |
| 15442.5 | 0.991019 |
| 19854.5 | 1.00036 |
| 24266.5 | 0.991 |
| 28678.5 | 1.04027 |
| 33090.5 | 1.05582 |
| 37502.5 | 1.03219 |
| 41914.5 | 1.0789 |
| 46326.5 | 1.04692 |
| 50738.5 | 1.06744 |
| 55150.5 | 1.06134 |
| 59562.5 | 1.09635 |
| 63974.5 | 1.20993 |
| 68386.5 | 1.07888 |
| 72798.5 | 1.12065 |
| 77210.5 | 1.0212 |
| 81622.5 | 1.07747 |
| 86034.5 | 1.0306 |
| 90446.5 | 0.947381 |
| 94858.5 | 0.989251 |
| 99270.5 | 0.947901 |
| 103682 | 1.13794 |
| 108094 | 1.14114 |
| 112506 | 1.13502 |
| 116918 | 1.08363 |
| 121330 | 1.02646 |
| 125742 | 1.04795 |
| 130154 | 0.971301 |
| 134566 | 1.06166 |
| 138978 | 1.14547 |
| 143390 | 1.08133 |
| 147802 | 1.06102 |
| 152214 | 0.981925 |
| 156626 | 1.06058 |
| 161038 | 1.01752 |
| 165450 | 0.965023 |
| 169862 | 0.96208 |
| 174274 | 1.02802 |
| 178686 | 0.960484 |
| 183098 | 0.976245 |
| 187510 | 0.93546 |
| 191922 | 1.04518 |
| 196334 | 1.09096 |
| 200746 | 0.99373 |
| 205158 | 1.05417 |
| 209570 | 1.07507 |
| 213982 | 1.07732 |
| 218394 | 1.07337 |
| 222806 | 1.07479 |
| 227218 | 1.03969 |
| 231630 | 1.00183 |
| 236042 | 0.924522 |
| 240454 | 0.985907 |
| 244866 | 0.942818 |
| 249278 | 1.01197 |
| 253690 | 1.02415 |
| 258102 | 1.08056 |
| 262514 | 0.944932 |
| 266926 | 0.967577 |
| 271338 | 1.03465 |
| 275750 | 0.936761 |
| 280162 | 0.947419 |
| 284574 | 0.915155 |
| 288986 | 0.967374 |
| 293398 | 0.992091 |
| 297810 | 1.10969 |
| 302222 | 1.11819 |
| 306634 | 0.965099 |
| 311046 | 0.97138 |
| 315458 | 0.985754 |
| 319870 | 0.920724 |
| 324282 | 0.9144 |
| 328694 | 1.01989 |
| 333106 | 0.927921 |
| 337518 | 0.646881 |
| 341930 | 0.897632 |
| 346342 | 0.967145 |
| 350754 | 0.941063 |
| 355166 | 0.846001 |
| 359578 | 0.962068 |
| 363990 | 0.925956 |
| 368402 | 0.900912 |
| 372814 | 0.942022 |
| 377226 | 1.08795 |
| 381638 | 1.13709 |
| 386050 | 1.05224 |
| 390462 | 1.06842 |
| 394874 | 1.03718 |
| 399286 | 1.04553 |
| 403698 | 0.973686 |
| 408110 | 1.00624 |
| 412522 | 1.07965 |
| 416934 | 1.07831 |
| 421346 | 1.03863 |
| 425758 | 0.801141 |
| 430170 | 0.978045 |
| 434582 | 0.868317 |
| 438994 | 0.978264 |
| 443406 | 0.908479 |
| 447818 | 1.0325 |
| 452230 | 0.980063 |
| 456642 | 0.850081 |
| 461054 | 0.938877 |
| 465466 | 0.893218 |
| 469878 | 0.924778 |
| 474290 | 0.969231 |
| 478702 | 1.02858 |
| 483114 | 1.16341 |
| 487526 | 1.11386 |
| 491938 | 1.00007 |
| 496350 | 0.909719 |
| 500762 | 0.927585 |
| 505174 | 1.06415 |
| 509586 | 0.957474 |
| 513998 | 0.949845 |
| 518410 | 0.968683 |
| 522822 | 0.964256 |
| 527234 | 0.98775 |
| 531646 | 0.884993 |
| 536058 | 0.960896 |
| 540470 | 1.01122 |
| 544882 | 1.0108 |
| 549294 | 1.0275 |
| 553706 | 1.0474 |
| 558118 | 1.04662 |
| 562530 | 1.07938 |
| 566942 | 0.980417 |
| 571354 | 0.905641 |
| 575766 | 0.941182 |
| 580178 | 0.988714 |
| 584590 | 0.933837 |
| 589002 | 0.986331 |
| 593414 | 1.02271 |
| 597826 | 1.08659 |
| 602238 | 1.0923 |
| 606650 | 1.07583 |
| 611062 | 0.94485 |
| 615474 | 0.883604 |
| 619886 | 0.935903 |
| 624298 | 0.743735 |
| 628710 | 0.844524 |
| 633122 | 0.832237 |
| 637534 | 0.9553 |
| 641946 | 0.969903 |
| 646358 | 0.995972 |
| 650770 | 1.13132 |
| 655182 | 1.10942 |
| 659594 | 1.00787 |
| 664006 | 1.05113 |
| 668418 | 0.953642 |
| 672830 | 0.740482 |
| 677242 | 0.931624 |
| 681654 | 1.08002 |
| 686066 | 1.14216 |
| 690478 | 1.05717 |
| 694890 | 1.06231 |
| 699302 | 0.933366 |
| 703714 | 0.962804 |
| 708126 | 0.961597 |
| 712538 | 0.865491 |
| 716950 | 0.938227 |
| 721362 | 0.963507 |
| 725774 | 0.881411 |
| 730186 | 0.969464 |
| 734598 | 0.974745 |
| 739010 | 1.00342 |
| 743422 | 0.872763 |
| 747834 | 0.773724 |
| 752246 | 0.946915 |
| 756658 | 1.04973 |
| 761070 | 1.2482 |
| 765482 | 1.21147 |
| 769894 | 0.999221 |
| 774306 | 1.01354 |
| 778718 | 1.05129 |
| 783130 | 1.10277 |
| 787542 | 1.02586 |
| 791954 | 0.993065 |
| 796366 | 1.0338 |
| 800778 | 1.09303 |
| 805190 | 0.926106 |
| 809602 | 0.968013 |
| 814014 | 0.957933 |
| 818426 | 0.945874 |
| 822838 | 0.895633 |
| 827250 | 1.00515 |
| 831662 | 0.838751 |
| 836074 | 0.779419 |
| 840486 | 0.873151 |
| 844898 | 0.917691 |
| 849310 | 0.924263 |
| 853722 | 0.996829 |
| 858134 | 0.9805 |
| 862546 | 0.938051 |
| 866958 | 0.978904 |
| 871370 | 0.904651 |
| 875782 | 1.10133 |
| 880194 | 1.02747 |

Coverage across reference - H37Rv - 102

|  | Norm |
| --- | --- |
| 2206.5 | 1.21756 |
| 6618.5 | 1.50186 |
| 11030.5 | 1.16389 |
| 15442.5 | 1.0255 |
| 19854.5 | 1.04329 |
| 24266.5 | 1.02065 |
| 28678.5 | 1.06054 |
| 33090.5 | 1.03766 |
| 37502.5 | 1.01819 |
| 41914.5 | 1.02157 |
| 46326.5 | 1.1005 |
| 50738.5 | 1.18608 |
| 55150.5 | 1.05313 |
| 59562.5 | 1.13232 |
| 63974.5 | 1.15501 |
| 68386.5 | 1.10894 |
| 72798.5 | 1.11173 |
| 77210.5 | 0.970275 |
| 81622.5 | 1.04915 |
| 86034.5 | 1.07946 |
| 90446.5 | 0.91507 |
| 94858.5 | 0.960083 |
| 99270.5 | 0.922247 |
| 103682 | 1.05639 |
| 108094 | 1.08904 |
| 112506 | 1.12983 |
| 116918 | 0.948125 |
| 121330 | 0.96912 |
| 125742 | 1.05904 |
| 130154 | 1.04773 |
| 134566 | 1.13512 |
| 138978 | 1.20076 |
| 143390 | 1.0885 |
| 147802 | 1.09839 |
| 152214 | 0.991226 |
| 156626 | 1.08763 |
| 161038 | 1.01047 |
| 165450 | 0.947391 |
| 169862 | 0.985695 |
| 174274 | 1.0748 |
| 178686 | 0.998194 |
| 183098 | 1.03501 |
| 187510 | 0.902827 |
| 191922 | 1.0803 |
| 196334 | 1.15087 |
| 200746 | 0.961118 |
| 205158 | 1.0562 |
| 209570 | 1.07458 |
| 213982 | 0.994626 |
| 218394 | 1.06412 |
| 222806 | 1.05358 |
| 227218 | 1.08398 |
| 231630 | 1.0791 |
| 236042 | 0.910146 |
| 240454 | 0.997854 |
| 244866 | 0.958442 |
| 249278 | 1.04528 |
| 253690 | 1.05775 |
| 258102 | 1.09183 |
| 262514 | 1.00176 |
| 266926 | 0.992239 |
| 271338 | 1.01566 |
| 275750 | 0.961962 |
| 280162 | 0.988415 |
| 284574 | 0.887371 |
| 288986 | 0.95952 |
| 293398 | 1.03431 |
| 297810 | 1.23685 |
| 302222 | 1.17581 |
| 306634 | 1.00585 |
| 311046 | 0.990049 |
| 315458 | 0.950417 |
| 319870 | 0.916207 |
| 324282 | 0.994484 |
| 328694 | 1.03257 |
| 333106 | 1.00593 |
| 337518 | 0.650718 |
| 341930 | 0.880122 |
| 346342 | 0.933184 |
| 350754 | 0.932808 |
| 355166 | 0.827238 |
| 359578 | 0.963026 |
| 363990 | 1.01912 |
| 368402 | 0.939483 |
| 372814 | 0.948699 |
| 377226 | 1.06812 |
| 381638 | 1.09635 |
| 386050 | 1.08247 |
| 390462 | 1.07466 |
| 394874 | 1.1013 |
| 399286 | 1.08548 |
| 403698 | 1.04512 |
| 408110 | 1.00676 |
| 412522 | 1.05166 |
| 416934 | 1.06789 |
| 421346 | 1.08898 |
| 425758 | 0.832875 |
| 430170 | 0.972355 |
| 434582 | 0.839704 |
| 438994 | 1.00909 |
| 443406 | 0.930958 |
| 447818 | 0.920982 |
| 452230 | 0.935319 |
| 456642 | 0.869703 |
| 461054 | 0.995938 |
| 465466 | 0.938745 |
| 469878 | 0.873812 |
| 474290 | 0.986119 |
| 478702 | 1.01908 |
| 483114 | 1.2212 |
| 487526 | 1.145 |
| 491938 | 1.07249 |
| 496350 | 0.965402 |
| 500762 | 1.03152 |
| 505174 | 1.02323 |
| 509586 | 0.865528 |
| 513998 | 0.84057 |
| 518410 | 0.917918 |
| 522822 | 0.983728 |
| 527234 | 0.998845 |
| 531646 | 0.920952 |
| 536058 | 0.962222 |
| 540470 | 0.966839 |
| 544882 | 1.08815 |
| 549294 | 1.08343 |
| 553706 | 1.0267 |
| 558118 | 1.07489 |
| 562530 | 1.10631 |
| 566942 | 1.00406 |
| 571354 | 0.882662 |
| 575766 | 0.926568 |
| 580178 | 1.05168 |
| 584590 | 0.871695 |
| 589002 | 1.03912 |
| 593414 | 0.982321 |
| 597826 | 1.0227 |
| 602238 | 1.0552 |
| 606650 | 1.0799 |
| 611062 | 0.988963 |
| 615474 | 0.874499 |
| 619886 | 0.871743 |
| 624298 | 0.794662 |
| 628710 | 0.932523 |
| 633122 | 0.808455 |
| 637534 | 1.00161 |
| 641946 | 0.919541 |
| 646358 | 0.957378 |
| 650770 | 1.1759 |
| 655182 | 1.06407 |
| 659594 | 1.01082 |
| 664006 | 1.02464 |
| 668418 | 0.898985 |
| 672830 | 0.676955 |
| 677242 | 0.954834 |
| 681654 | 1.13448 |
| 686066 | 1.24054 |
| 690478 | 1.06603 |
| 694890 | 1.05748 |
| 699302 | 0.914782 |
| 703714 | 0.957276 |
| 708126 | 0.959904 |
| 712538 | 0.910859 |
| 716950 | 0.910384 |
| 721362 | 0.974761 |
| 725774 | 0.869674 |
| 730186 | 0.954709 |
| 734598 | 0.942031 |
| 739010 | 1.05265 |
| 743422 | 0.895527 |
| 747834 | 0.825527 |
| 752246 | 0.950374 |
| 756658 | 1.08858 |
| 761070 | 1.2877 |
| 765482 | 1.17966 |
| 769894 | 1.00597 |
| 774306 | 0.979338 |
| 778718 | 1.05957 |
| 783130 | 1.1537 |
| 787542 | 1.1001 |
| 791954 | 1.06606 |
| 796366 | 1.07234 |
| 800778 | 1.09033 |
| 805190 | 1.00866 |
| 809602 | 0.937356 |
| 814014 | 1.01952 |
| 818426 | 0.972973 |
| 822838 | 0.844559 |
| 827250 | 1.00172 |
| 831662 | 0.909342 |
| 836074 | 0.784525 |
| 840486 | 0.885298 |
| 844898 | 0.894134 |
| 849310 | 0.863726 |
| 853722 | 0.990144 |
| 858134 | 1.04939 |
| 862546 | 1.00018 |
| 866958 | 0.986393 |
| 871370 | 0.855274 |
| 875782 | 1.05855 |
| 880194 | 1.07057 |

Genomic window

Supplementary Figure 4: Normalized sequencing depth across the reference genome using different input amounts of target DNA for all H37Rv LoD samples. Values represent the average sequencing depth across 1,000 bp windows normalized by the overall median sequencing depth of the sample.

B

|  | Normalised |
| --- | --- |
| 2206.5 | 0.925977 |
| 6618.5 | 1.27106 |
| 11030.5 | 1.10459 |
| 15442.5 | 0.901086 |
| 19854.5 | 0.976418 |
| 24266.5 | 0.974038 |
| 28678.5 | 1.08305 |
| 33090.5 | 1.0177 |
| 37502.5 | 0.988317 |
| 41914.5 | 1.05891 |
| 46326.5 | 0.977922 |
| 50738.5 | 0.987884 |
| 55150.5 | 1.03993 |
| 59562.5 | 0.94687 |
| 63974.5 | 0.933343 |
| 68386.5 | 1.00988 |
| 72798.5 | 1.04124 |
| 77210.5 | 0.996044 |
| 81622.5 | 0.481353 |
| 86034.5 | 1.02458 |
| 90446.5 | 0.931509 |
| 94858.5 | 0.963581 |
| 99270.5 | 0.908122 |
| 103682 | 1.07758 |
| 108094 | 0.998012 |
| 112506 | 0.923216 |
| 116918 | 0.889135 |
| 121330 | 0.957307 |
| 125742 | 0.865563 |
| 130154 | 1.08331 |
| 134566 | 0.946674 |
| 138978 | 0.940555 |
| 143390 | 0.9295 |
| 147802 | 1.13676 |
| 152214 | 0.962252 |
| 156626 | 1.05951 |
| 161038 | 1.24124 |
| 165450 | 0.950734 |
| 169862 | 0.97615 |
| 174274 | 0.956627 |
| 178686 | 0.978468 |
| 183098 | 0.9329 |
| 187510 | 0.875546 |
| 191922 | 0.965476 |
| 196334 | 0.932653 |
| 200746 | 0.942759 |
| 205158 | 0.979436 |
| 209570 | 1.06071 |
| 213982 | 0.928727 |
| 218394 | 0.982733 |
| 222806 | 1.03887 |
| 227218 | 0.965033 |
| 231630 | 1.10764 |
| 236042 | 1.24076 |
| 240454 | 0.968588 |
| 244866 | 0.985391 |
| 249278 | 1.02038 |
| 253690 | 1.18768 |
| 258102 | 1.01729 |
| 262514 | 0.946396 |
| 266926 | 0.880368 |
| 271338 | 1.04985 |
| 275750 | 0.919496 |
| 280162 | 1.2891 |
| 284574 | 1.14272 |
| 288986 | 1.07665 |
| 293398 | 0.927409 |
| 297810 | 0.921093 |
| 302222 | 0.904723 |
| 306634 | 1.02867 |
| 311046 | 0.944913 |
| 315458 | 0.969762 |
| 319870 | 0.961057 |
| 324282 | 0.965188 |
| 328694 | 0.893689 |
| 333106 | 1.23193 |
| 337518 | 0.703258 |
| 341930 | 0.931004 |
| 346342 | 0.902765 |
| 350754 | 1.03347 |
| 355166 | 0.881006 |
| 359578 | 0.894204 |
| 363990 | 0.876391 |
| 368402 | 1.10333 |
| 372814 | 1.0419 |
| 377226 | 0.990141 |
| 381638 | 1.12983 |
| 386050 | 0.993097 |
| 390462 | 0.955823 |
| 394874 | 0.964281 |
| 399286 | 1.18436 |
| 403698 | 1.00061 |
| 408110 | 1.06142 |
| 412522 | 1.01701 |
| 416934 | 0.994128 |
| 421346 | 1.03841 |
| 425758 | 0.948519 |
| 430170 | 0.978561 |
| 434582 | 0.858578 |
| 438994 | 0.994282 |
| 443406 | 0.94583 |
| 447818 | 1.04908 |
| 452230 | 0.881655 |
| 456642 | 0.961118 |
| 461054 | 1.11975 |
| 465466 | 0.927811 |
| 469878 | 0.943099 |
| 474290 | 1.103 |
| 478702 | 0.97003 |
| 483114 | 0.966692 |
| 487526 | 0.964158 |
| 491938 | 0.985237 |
| 496350 | 1.01326 |
| 500762 | 0.948055 |
| 505174 | 1.05166 |
| 509586 | 1.02149 |
| 513998 | 1.0103 |
| 518410 | 1.11906 |
| 522822 | 1.0418 |
| 527234 | 1.06828 |
| 531646 | 1.12401 |
| 536058 | 0.942296 |
| 540470 | 0.939246 |
| 544882 | 1.05579 |
| 549294 | 0.947715 |
| 553706 | 0.999052 |
| 558118 | 1.28217 |
| 562530 | 1.30023 |
| 566942 | 1.09089 |
| 571354 | 0.944707 |
| 575766 | 0.971029 |
| 580178 | 1.04449 |
| 584590 | 1.03067 |
| 589002 | 1.1979 |
| 593414 | 0.889516 |
| 597826 | 1.06443 |
| 602238 | 0.975521 |
| 606650 | 1.06813 |
| 611062 | 0.942636 |
| 615474 | 0.944944 |
| 619886 | 0.885653 |
| 624298 | 0.975068 |
| 628710 | 0.890856 |
| 633122 | 1.05567 |
| 637534 | 0.976346 |
| 641946 | 0.908916 |
| 646358 | 0.985607 |
| 650770 | 1.04712 |
| 655182 | 0.98807 |
| 659594 | 1.07429 |
| 664006 | 1.30117 |
| 668418 | 0.832183 |
| 672830 | 0.920506 |
| 677242 | 0.894049 |
| 681654 | 0.879976 |
| 686066 | 0.953103 |
| 690478 | 0.916838 |
| 694890 | 1.04086 |
| 699302 | 1.04106 |
| 703714 | 1.06617 |
| 708126 | 0.997012 |
| 712538 | 1.03394 |
| 716950 | 1.05542 |
| 721362 | 1.05779 |
| 725774 | 1.09844 |
| 730186 | 0.962468 |
| 734598 | 1.32519 |
| 739010 | 1.1589 |
| 743422 | 1.22908 |
| 747834 | 0.938731 |
| 752246 | 1.00145 |
| 756658 | 1.26593 |
| 761070 | 1.20646 |
| 765482 | 1.52368 |
| 769894 | 1.0984 |
| 774306 | 1.00604 |
| 778718 | 1.11026 |
| 783130 | 1.23261 |
| 787542 | 1.25256 |
| 791954 | 0.998805 |
| 796366 | 0.966445 |
| 800778 | 1.13855 |
| 805190 | 1.17767 |
| 809602 | 0.956214 |
| 814014 | 1.16141 |
| 818426 | 1.0167 |
| 822838 | 0.919321 |
| 827250 | 1.11342 |
| 831662 | 1.16957 |
| 836074 | 0.959068 |
| 840486 | 1.24675 |
| 844898 | 0.798586 |
| 849310 | 1.00279 |
| 853722 | 0.918167 |
| 858134 | 0.934044 |
| 862546 | 0.983928 |
| 866958 | 0.888517 |
| 871370 | 0.902611 |
| 875782 | 1.02175 |
| 880194 | 0.948519 |

D

|  | Norm |
| --- | --- |
| 2206.5 | 0.86609 |
| 6618.5 | 1.40017 |
| 11030.5 | 1.16622 |
| 15442.5 | 1.09287 |
| 19854.5 | 1.09505 |
| 24266.5 | 1.1025 |
| 28678.5 | 1.07781 |
| 33090.5 | 1.02157 |
| 37502.5 | 1.22614 |
| 41914.5 | 1.14257 |
| 46326.5 | 0.995254 |
| 50738.5 | 1.09093 |
| 55150.5 | 1.24661 |
| 59562.5 | 1.11204 |
| 63974.5 | 1.26596 |
| 68386.5 | 0.925912 |
| 72798.5 | 1.15019 |
| 77210.5 | 1.05165 |
| 81622.5 | 0.475748 |
| 86034.5 | 1.12601 |
| 90446.5 | 1.02122 |
| 94858.5 | 1.03177 |
| 99270.5 | 0.839302 |
| 103682 | 1.05631 |
| 108094 | 1.08502 |
| 112506 | 1.06706 |
| 116918 | 1.19193 |
| 121330 | 0.981641 |
| 125742 | 1.06127 |
| 130154 | 1.00038 |
| 134566 | 0.885851 |
| 138978 | 0.990523 |
| 143390 | 1.0351 |
| 147802 | 1.07719 |
| 152214 | 1.04408 |
| 156626 | 1.09612 |
| 161038 | 1.22807 |
| 165450 | 0.977887 |
| 169862 | 0.963395 |
| 174274 | 1.06268 |
| 178686 | 0.864857 |
| 183098 | 1.06053 |
| 187510 | 0.788687 |
| 191922 | 1.01339 |
| 196334 | 1.07439 |
| 200746 | 0.952955 |
| 205158 | 0.996586 |
| 209570 | 1.06858 |
| 213982 | 1.06971 |
| 218394 | 0.976187 |
| 222806 | 0.976456 |
| 227218 | 1.06023 |
| 231630 | 1.06462 |
| 236042 | 1.01244 |
| 240454 | 1.1419 |
| 244866 | 0.898841 |
| 249278 | 1.05934 |
| 253690 | 1.01595 |
| 258102 | 1.1686 |
| 262514 | 1.0672 |
| 266926 | 1.01296 |
| 271338 | 0.992039 |
| 275750 | 1.0595 |
| 280162 | 0.996671 |
| 284574 | 0.963424 |
| 288986 | 0.950958 |
| 293398 | 1.0781 |
| 297810 | 0.964104 |
| 302222 | 1.15238 |
| 306634 | 0.993399 |
| 311046 | 1.03573 |
| 315458 | 1.13054 |
| 319870 | 0.975323 |
| 324282 | 1.03465 |
| 328694 | 1.07545 |
| 333106 | 0.953012 |
| 337518 | 0.454867 |
| 341930 | 0.867265 |
| 346342 | 1.1593 |
| 350754 | 1.02122 |
| 355166 | 0.934964 |
| 359578 | 0.786959 |
| 363990 | 0.745793 |
| 368402 | 0.781802 |
| 372814 | 0.837517 |
| 377226 | 0.966852 |
| 381638 | 1.12904 |
| 386050 | 1.13125 |
| 390462 | 0.983879 |
| 394874 | 1.05887 |
| 399286 | 1.06171 |
| 403698 | 1.11596 |
| 408110 | 1.10993 |
| 412522 | 1.10976 |
| 416934 | 1.09454 |
| 421346 | 1.0469 |
| 425758 | 0.805417 |
| 430170 | 0.897807 |
| 434582 | 0.782822 |
| 438994 | 1.08691 |
| 443406 | 1.02038 |
| 447818 | 0.975436 |
| 452230 | 1.15741 |
| 456642 | 0.801734 |
| 461054 | 1.05213 |
| 465466 | 0.873017 |
| 469878 | 0.955009 |
| 474290 | 1.03228 |
| 478702 | 1.01303 |
| 483114 | 1.00891 |
| 487526 | 0.97759 |
| 491938 | 1.05265 |
| 496350 | 1.02114 |
| 500762 | 1.03872 |
| 505174 | 1.15925 |
| 509586 | 1.12652 |
| 513998 | 1.08546 |
| 518410 | 1.0187 |
| 522822 | 0.930573 |
| 527234 | 1.19436 |
| 531646 | 1.18679 |
| 536058 | 0.951737 |
| 540470 | 0.985324 |
| 544882 | 0.98208 |
| 549294 | 1.01657 |
| 553706 | 1.11045 |
| 558118 | 1.00051 |
| 562530 | 1.09587 |
| 566942 | 1.06985 |
| 571354 | 0.981825 |
| 575766 | 0.888543 |
| 580178 | 1.00201 |
| 584590 | 1.01703 |
| 589002 | 1.15845 |
| 593414 | 0.968608 |
| 597826 | 0.944668 |
| 602238 | 1.23387 |
| 606650 | 1.15643 |
| 611062 | 0.88435 |
| 615474 | 0.980777 |
| 619886 | 0.926819 |
| 624298 | 0.795161 |
| 628710 | 0.864588 |
| 633122 | 0.947614 |
| 637534 | 1.0249 |
| 641946 | 1.08654 |
| 646358 | 0.990254 |
| 650770 | 1.01262 |
| 655182 | 1.05348 |
| 659594 | 1.03626 |
| 664006 | 1.02393 |
| 668418 | 0.964033 |
| 672830 | 0.700547 |
| 677242 | 0.93114 |
| 681654 | 1.02343 |
| 686066 | 1.02091 |
| 690478 | 0.906222 |
| 694890 | 1.04979 |
| 699302 | 0.953592 |
| 703714 | 0.890696 |
| 708126 | 0.992804 |
| 712538 | 0.935063 |
| 716950 | 0.927782 |
| 721362 | 1.13067 |
| 725774 | 0.917002 |
| 730186 | 1.0848 |
| 734598 | 1.0139 |
| 739010 | 1.09024 |
| 743422 | 0.902879 |
| 747834 | 0.878272 |
| 752246 | 1.03186 |
| 756658 | 1.0469 |
| 761070 | 1.45898 |
| 765482 | 1.41039 |
| 769894 | 1.0476 |
| 774306 | 1.01077 |
| 778718 | 1.06872 |
| 783130 | 1.46211 |
| 787542 | 1.06424 |
| 791954 | 0.994348 |
| 796366 | 0.987081 |
| 800778 | 1.1569 |
| 805190 | 1.07062 |
| 809602 | 1.11168 |
| 814014 | 1.08811 |
| 818426 | 1.08552 |
| 822838 | 0.873867 |
| 827250 | 1.12972 |
| 831662 | 0.733667 |
| 836074 | 0.709046 |
| 840486 | 0.854445 |
| 844898 | 0.814937 |
| 849310 | 0.805885 |
| 853722 | 0.917512 |
| 858134 | 1.06658 |
| 862546 | 1.0121 |
| 866958 | 0.993229 |
| 871370 | 0.948294 |
| 875782 | 1.14288 |
| 880194 | 1.02449 |

Normalised sequencing depth

W

|  | Norm |
| --- | --- |
| 2206.5 | 0.672201 |
| 6618.5 | 1.46646 |
| 11030.5 | 1.2453 |
| 15442.5 | 1.19917 |
| 19854.5 | 1.20756 |
| 24266.5 | 0.975748 |
| 28678.5 | 1.02658 |
| 33090.5 | 1.14574 |
| 37502.5 | 0.973736 |
| 41914.5 | 1.28267 |
| 46326.5 | 1.20172 |
| 50738.5 | 1.29431 |
| 55150.5 | 0.992492 |
| 59562.5 | 1.11455 |
| 63974.5 | 1.24493 |
| 68386.5 | 1.04015 |
| 72798.5 | 1.22609 |
| 77210.5 | 0.978468 |
| 81622.5 | 0.384038 |
| 86034.5 | 1.03595 |
| 90446.5 | 1.21575 |
| 94858.5 | 0.997988 |
| 99270.5 | 0.99643 |
| 103682 | 0.986146 |
| 108094 | 1.2066 |
| 112506 | 1.03006 |
| 116918 | 1.17345 |
| 121330 | 1.08199 |
| 125742 | 1.0757 |
| 130154 | 0.839075 |
| 134566 | 1.17226 |
| 138978 | 1.05261 |
| 143390 | 1.18645 |
| 147802 | 1.07448 |
| 152214 | 1.10656 |
| 156626 | 1.17319 |
| 161038 | 1.08658 |
| 165450 | 1.08536 |
| 169862 | 1.08143 |
| 174274 | 1.09905 |
| 178686 | 0.937188 |
| 183098 | 0.918631 |
| 187510 | 0.909168 |
| 191922 | 0.995099 |
| 196334 | 1.01371 |
| 200746 | 1.07675 |
| 205158 | 1.11358 |
| 209570 | 1.166 |
| 213982 | 1.05312 |
| 218394 | 1.08879 |
| 222806 | 0.969883 |
| 227218 | 1.12517 |
| 231630 | 1.13914 |
| 236042 | 0.979601 |
| 240454 | 1.05247 |
| 244866 | 1.0997 |
| 249278 | 1.16194 |
| 253690 | 1.04777 |
| 258102 | 1.1131 |
| 262514 | 1.06703 |
| 266926 | 1.11253 |
| 271338 | 1.0064 |
| 275750 | 0.936933 |
| 280162 | 1.14112 |
| 284574 | 1.11996 |
| 288986 | 1.18175 |
| 293398 | 1.10661 |
| 297810 | 1.16146 |
| 302222 | 1.18053 |
| 306634 | 0.922257 |
| 311046 | 1.00391 |
| 315458 | 0.978383 |
| 319870 | 0.874207 |
| 324282 | 1.17484 |
| 328694 | 1.11197 |
| 333106 | 0.933619 |
| 337518 | 0.300544 |
| 341930 | 0.78431 |
| 346342 | 1.06219 |
| 350754 | 0.964614 |
| 355166 | 0.888911 |
| 359578 | 0.851201 |
| 363990 | 0.75408 |
| 368402 | 0.670501 |
| 372814 | 0.861854 |
| 377226 | 0.896929 |
| 381638 | 1.02961 |
| 386050 | 1.12605 |
| 390462 | 0.956284 |
| 394874 | 1.06253 |
| 399286 | 1.12169 |
| 403698 | 1.06958 |
| 408110 | 1.09876 |
| 412522 | 1.13698 |
| 416934 | 1.21932 |
| 421346 | 1.11596 |
| 425758 | 0.833607 |
| 430170 | 0.925714 |
| 434582 | 0.900244 |
| 438994 | 1.1828 |
| 443406 | 0.92849 |
| 447818 | 0.997082 |
| 452230 | 1.40095 |
| 456642 | 1.08868 |
| 461054 | 1.07273 |
| 465466 | 1.16155 |
| 469878 | 1.02068 |
| 474290 | 0.897779 |
| 478702 | 0.947926 |
| 483114 | 0.877522 |
| 487526 | 0.897807 |
| 491938 | 1.06188 |
| 496350 | 0.947246 |
| 500762 | 0.972348 |
| 505174 | 1.16699 |
| 509586 | 1.0236 |
| 513998 | 1.0808 |
| 518410 | 1.22861 |
| 522822 | 1.13171 |
| 527234 | 1.15602 |
| 531646 | 1.2292 |
| 536058 | 0.942571 |
| 540470 | 0.942628 |
| 544882 | 0.986117 |
| 549294 | 1.00082 |
| 553706 | 1.06131 |
| 558118 | 1.0844 |
| 562530 | 1.10826 |
| 566942 | 0.902539 |
| 571354 | 1.00561 |
| 575766 | 0.988441 |
| 580178 | 1.243 |
| 584590 | 1.06681 |
| 589002 | 0.875737 |
| 593414 | 1.06924 |
| 597826 | 1.00725 |
| 602238 | 0.965803 |
| 606650 | 1.22858 |
| 611062 | 1.12684 |
| 615474 | 1.00017 |
| 619886 | 1.16013 |
| 624298 | 0.892452 |
| 628710 | 0.944668 |
| 633122 | 1.02289 |
| 637534 | 1.32162 |
| 641946 | 1.01224 |
| 646358 | 1.06754 |
| 650770 | 1.25994 |
| 655182 | 1.04882 |
| 659594 | 0.974473 |
| 664006 | 1.165 |
| 668418 | 0.843098 |
| 672830 | 0.728808 |
| 677242 | 0.911916 |
| 681654 | 0.912795 |
| 686066 | 1.07375 |
| 690478 | 1.06052 |
| 694890 | 1.17053 |
| 699302 | 0.980224 |
| 703714 | 1.04281 |
| 708126 | 0.953734 |
| 712538 | 0.918574 |
| 716950 | 0.927669 |
| 721362 | 1.14914 |
| 725774 | 0.927414 |
| 730186 | 0.927754 |
| 734598 | 1.375 |
| 739010 | 1.32318 |
| 743422 | 0.961214 |
| 747834 | 0.848906 |
| 752246 | 1.03184 |
| 756658 | 0.980819 |
| 761070 | 1.52833 |
| 765482 | 1.39058 |
| 769894 | 1.11528 |
| 774306 | 1.10412 |
| 778718 | 1.24878 |
| 783130 | 1.30715 |
| 787542 | 1.07437 |
| 791954 | 1.01504 |
| 796366 | 0.966965 |
| 800778 | 1.20719 |
| 805190 | 1.11775 |
| 809602 | 1.11166 |
| 814014 | 1.142 |
| 818426 | 1.28842 |
| 822838 | 0.996345 |
| 827250 | 1.0389 |
| 831662 | 0.774025 |
| 836074 | 0.663928 |
| 840486 | 0.884208 |
| 844898 | 0.737817 |
| 849310 | 0.764223 |
| 853722 | 1.10571 |
| 858134 | 1.02136 |
| 862546 | 0.838055 |
| 866958 | 1.07012 |
| 871370 | 0.936197 |
| 875782 | 1.08692 |
| 880194 | 1.12579 |

Genomic window

Supplementary Figure 5: Normalized sequencing depth across the reference genome for all enriched sediment samples (B – Benzonase treated, D – DNase treated and W – Washed). Values represent the average sequencing depth across 1,000 bp windows normalized by the overall median sequencing depth of the sample.
